# ANCHOR : Atlas of Neurochemical Characterization of the Human Brainstem with 3D Reconstruction

**DOI:** 10.64898/2026.06.03.727794

**Authors:** Mihail Bota, Soundharya Venkatesh, Shevani Arun Arunesh, Nagajothi Ganesan, Supriti Mulay, Karan Ramana Gopi, Sruti Rekha Muni, Shrimathi Mani, Sam C. Chrisline, A.S.T. Aditya Bharg, Kanna G Vinoth, Rajeswaran Rangaswami, S. Lata, E. Harish Kumar, S. Suresh, Mousumi Sen, Ranjit Immanuel James, Abi Manesh, George M. Varghese, K.V. Vinoth, Keerthi Ram, Richa Verma, Paul R. Manger, Mohanasankar Sivaprakasam

## Abstract

The human brainstem is a complex division of the brain comprised of more than 200 nuclei and fiber tracts. The brainstem is essential for the functioning of the entire body. We introduce here the most detailed human brainstem Atlas across the human lifespan: fetus, child, adult. ANCHOR, the Atlas of Neurochemical Characterization of the Human Brainstem, is an online platform that includes more than 800 serial histological sections, stained for Nissl and seven immunochemical (IHC) markers, from the human brainstem of three ages: 25 fetal gestational weeks (GW), 9 years old, and 54 years old. This makes ANCHOR the most comprehensive human brainstem Atlas to date. In these three brainstems, we identified and manually annotated over 200 structures. We further characterized these structures with the seven IHC markers. We specifically describe the catecholaminergic groups in the human brainstem across all three age groups. In addition, we identified the protoplasmic commissural dendrites of the hypoglossal nucleus and we describe the pretectal nuclei in the Nissl-stained fetal 25 GW brainstem. ANCHOR includes an online viewer that integrates multimodal data, from magnetic resonance imaging and block face imaging to Nissl- and IHC-stained serial sections and 3D reconstruction of the entire brainstem. For the 9-year-old specimen, the online viewer allows simultaneous navigation of annotated sections with corresponding IHC, for viewing the specific region-wise cellular features accessible at https://anchor.humanbrain.in/.

## 1. Introduction

The brainstem is crucial for a wide range of brain functions and is the controller of the autonomic and enteric divisions of the nervous system. The brainstem forms the conduit between the cerebrum and most of the cranial nerves and spinal cord. In humans, the brainstem occupies less than 3% of the encephalic volume (Erbagci et al., 2012; Coulombe et al., 2021), compared to monkeys where it occupies about 4% (Scott et al., 2016), and 15% in rodents (Badea et al., 2007). The brainstem is comprised of more than 200 distinct structures, including nuclei and their component parts (Mezias et al., 2024). The mammalian brainstem includes many neurochemically distinct neuronal nuclei, such as the catecholaminergic (Dahlström & Fuxe, 1964; Pearson et al., 1983; Smeeets & Gonzalez, 2000; Manger & Eschenko, 2021), serotonergic (Hornung, 2003), and cholinergic (Mesulam et al., 1984; Mesulam et al., 1989) neuronal populations involved in modulation of functions across the entire brain.

The brainstem can be divided in four major divisions: midbrain, pons, medulla, and sometimes the cerebellum (Swanson, 2015), or into several neuromeres (Watson et al., 2019); however, many nuclei cross these neuromeric boundaries and the main brainstem divisions. Functionally, the brainstem can be divided based on dorsoventral and rostrocaudal axes (Nieuwenhuys et al., 2008). Thus, the brainstem is comprised of a dorsal sensory part, a motor ventral part, a reticular core, and the pre- and postcerebellar nuclei (Swanson, 1998, Nieuwenhuys et al., 2008, Manger, 2017). Finally, based on developmental gene expression patters, the caudal pons and medulla were included in the hindbrain, and the rostral part of the pons was renamed as isthmus (Paxinos, 2019; Watson et al., 2019). Herein, we use the traditional subdivisions of the brainstem (midbrain, pons, and medulla oblongata) for ease of description and comparison to earlier studies and studies in other species.

Even though the brainstem was recognized as a relay between the spinal cord, and the cerebrum by Willis as early as XVIIth century (Coulombe et al. 2021), it is not yet been described in as much detail as other major divisions of the brain. This may be due the large number of nuclei (and their parts) and fiber tracts that originate, end, or pass through it. There is also a lack of systematic large scale multimodal studies of the brainstem, and existing studies employ different neuroanatomical nomenclatures. In addition, the sectioning angles used in different studies vary, leading to difficulties in comparing different results. However, a detailed understanding of the chemical neuroanatomy of the human brainstem is important for understanding the cell types and pathways affected in brainstem lesions, or in diseases such as those occurring in sudden infant death syndrome (SIDS; Kinney et al., 1992).

The brainstem begins to develop from early gestational stages (Bayer & Altman, 2004-2006) originating from mesomeres 1 and 2, and rhombomeres 1-3, plus 5-11 (Paxinos et al., 2019). The development of the hindbrain in other vertebrate species is regularly studied due to availability of specimens (Bayer & Altman, 2004-2006; Puelles, 2018), while in humans there are only few detailed and comprehensive studies in the embryo (Müller & O’Rahilly, 2011), and fetus (Bayer & Altman, 2004-2006), respectively. To our knowledge, there are only two systematic, publicly available, cytoarchitectonic atlases that include the brainstem in different stages of fetal development: (1) The Allen Brain Atlas (AIBS) of the human fetus at 21 postconceptional weeks (pcw; Ding et al., 2017); and (2) DHARANI, that includes five cytoarchitectonic maps of the human fetal brain during the second trimester of development from 14-24 gestational weeks (GW; Verma et al., 2025).

For the adult human brainstem, the cytoarchitectonic (Nissl) atlas produced by Olszewski and Baxter (1954) remains a seminal work. The atlas produced by Paxinos and colleagues (2012; 2019; 2023) includes detailed cytoarchitectonic (Nissl) and acetylcholinesterase (AchE) sections and maps, as well as sections labeled for myelin, that are complemented by magnetic resonance imaging (MRI) scans. The publicly available AIBS human brain atlas includes parcellation maps of the adult human brainstem based on Nissl, parvalbumin (PV), and SMI-32 stains (Ding et al., 2017). Other efforts to parcellate the human adult brainstem, using Nissl and immunohistochemical (IHC) stains, include a topographic cytoarchitectonic and myeloarchitectonic atlas sectioned in the plane of the ponto-mesencephalic junction (Coulombe et al., 2021), and a cytoarchitectonic and IHC atlas, paired with 7 Tesla MRI imaging (Agostinelli et al., 2023); however, the former lacks IHC staining in the same specimen, and the latter includes a limited set of sections. Finally, the human brainstem has been investigated with MRI (Naidich et al., 2007; Rushmore et al., 2020; Adil et al., 2021), but the numbers of identifiable parts are limited due to the resolution of MRI.

An integrated approach for understanding the nuclei of the human brainstem needs to include their development, detailed multimodal maps, and volumetric (3D) reconstructions of the nuclei, as well as the fiber tracts. Moreover, it is preferable to have the data sets and maps publicly available. In this paper we present and briefly describe the Atlas of Neurochemical Characterization of the Human Brainstem (ANCHOR; https://anchor.humanbrain.in/), an online and publicly available platform that includes human brainstem Nissl staining, neuronal nuclei (NeuN), neurofilament heavy chain (NFH), tyrosine hydroxylase (TH), orexin-A (OX-A), calretinin (CR), calbindin (CB), parvalbumin (PV) and myelin IHC datasets across three ages: (1) the third trimester of gestation (25 GW); (2) 9 years of age; and (3) 54 years of age. The maps delineate more than 200 brainstem nuclei and fiber tracts, and were reconstructed from more than 800 serial sections. ANCHOR includes more than 200 Nissl and IHC manually annotated sections, MRI scans, block face imaging (BFI; Karthik et al., 2023), and 3D reconstructions of entire brainstems across for the three ages. ANCHOR serves not only to understand the complex architecture of the brainstem, but also to inform clinicians about its detailed organization. We also describe the pretectum in the human fetus (25 GW) as it is scarcely described in the literature (Verma et al., 2025).

## 2. Methods

ANCHOR was developed using the human brain histological pipeline and the computational platform established at the Sudha Gopalakrishnan Brain Centre (SGBC), IIT Madras, as described in detail in Verma et al., (2025). The entire pipeline of ANCHOR is shown in Figure 1.

**Figure 1.**
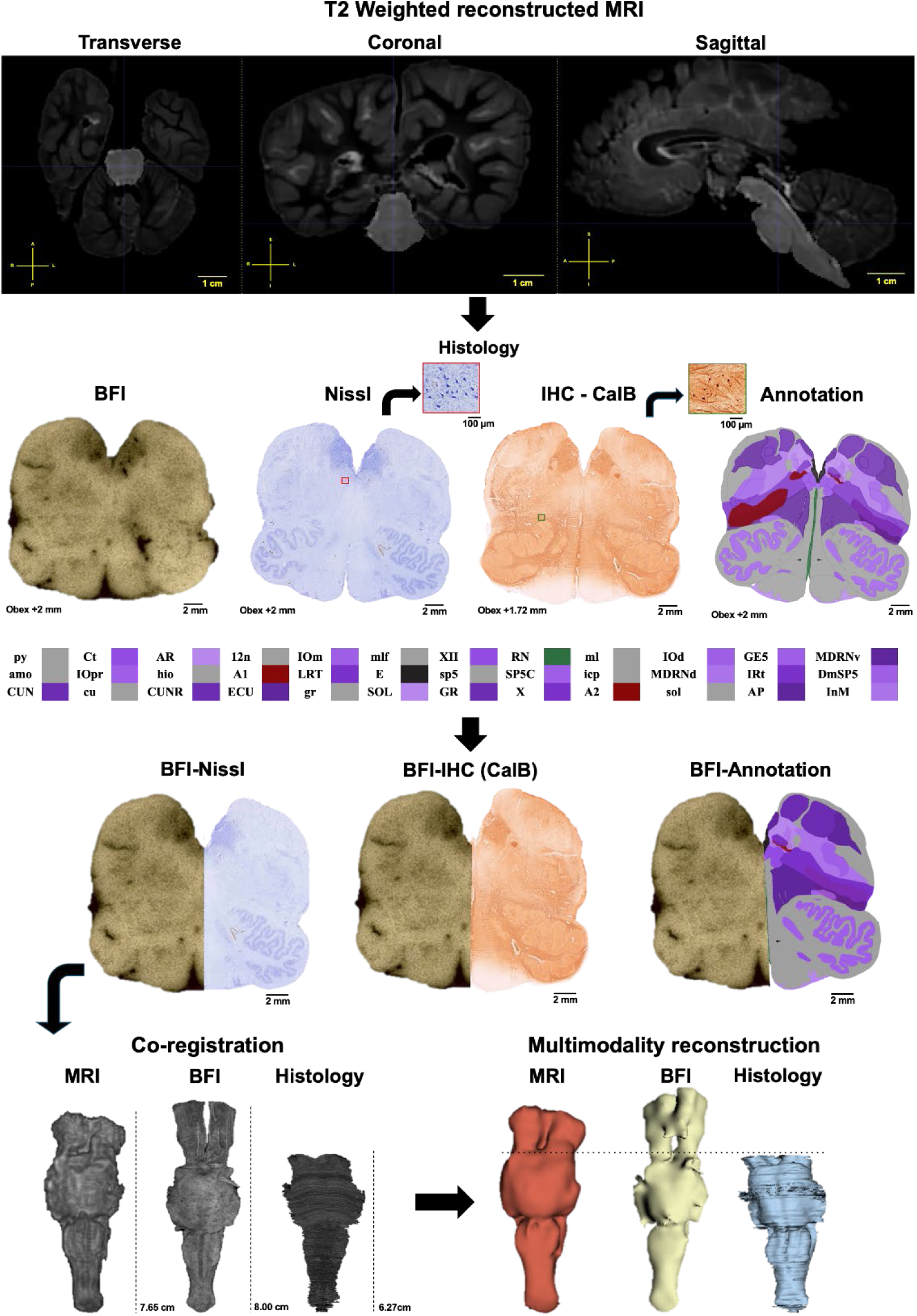
The ANCHOR pipeline for processing and 3D reconstruction.

### 2.1 Specimen details

Three specimens (S1 - 25GW, S2 - 9 years and S3 - 54 years) were acquired with due consent from the next of kin from Department of Pathology, Mediscan Systems Pvt. Ltd., Chennai, India (Mediscan), Department of Forensic Medicine, Government Kilpauk Medical College, Chennai, India, and Department of Infectious Diseases, Christian Medical College, Vellore, India. respectively. No known neurological or psychiatric disorders based on clinical and pathological assessment were identified. MRI for all three specimens were obtained from the Department of Radiology, Sri Ramachandra Institute of Higher Education and Research. Histological processing, digitization, quality control (QC), and atlas annotation followed the methodology described in Verma et al. (2025).

**Table 1.** Specimen details and parameters used in the MRI scanning.

| Specimen ID | Postmortem interval | MRI scanning interval | Plane of sectioning | Section thickness |
| --- | --- | --- | --- | --- |
| S1 | 2-4 hours | after 4 days of fixation | Transverse<br>(Coronal) | 20 μm |
| S2 | 19 hours and 50 minutes | after 14 days of fixation | Transverse along the longitudinal axis of the brainstem | 40 μm |
| S3 | 1 day and 21 hours | after 16 months of fixation | Transverse along the longitudinal axis of the brainstem | 20 μm |

### 2.2 MRI Acquisition

Ex vivo MRI data was acquired using a 3T GE SIGNA Architect scanner (see Table 2).

**Table 2.** The parameters used in MRI scan of the three specimens.

| Specimen | Sequence | Field of view (FOV)<br>(mm) | Spacing (mm) | Flip angle (degrees) | Echo time (TE)/Relaxation time (TR) (ms) | Slice thickness (mm) |
| --- | --- | --- | --- | --- | --- | --- |
| S1 | 3D Sagittal T1 MP RAGE | 512x512x140 | 0.293x0.293x0.5 | 8 | 2.36/2530.69 | 1 |
| S2 | Ax T2 FSE | 256x256x60 | 0.93x0.93x2 | 111 | 83.4/8018 | 2 |
|  | SAG T2 FSE | 256x256x75 | 0.8594x0.8594x2 | 111 | 84.84/10019 | 2 |
| S3 | 3D Sagittal T1 MP RAGE | 512x512x208 | 0.4492x0.4492x0.8 | 8 | 2.78/2528.36 | 1.6 |

The brainstem was considered as the region of interest and was manually segmented and delineated in ITK-SNAP (v4.4.0), guided by the Paxinos et al. (2019).

### 2.3 Histology processing

All three specimens were fixed in 4% paraformaldehyde in 0.01 M phosphate buffer (PB) at room temperature and subsequently cryoprotected with graded sucrose solutions (10%, 20%, and 30%) in 0.01 M PB at 4°C to minimize freezing artifacts. Prior to cryoprotection, 3D surface scanning of each brain was performed to generate a custom cryomold that was utilized to freeze the specimens in OCT medium at −80°C. The cryoblocks were stored at −80°C until sectioning, which was carried out using the tape-transfer technique. To establish a 3D reference volume, the exposed block face was imaged serially using a Block Face Imaging (BFI) camera. Specimen 1 was sectioned coronally as a whole brain at 20 μm thickness, whereas for specimens 2 and 3 the brainstem was dissected from the whole brain and sectioned in a transverse plane relative to its longitudinal axis at 40 μm and 20 μm thickness, respectively. Sections from specimen 1 were divided into three series and stained for Nissl, hematoxylin and eosin (H&E), and either immunohistochemistry (IHC) or myelin. For specimens 2 and 3, the sections were divided into ten series and stained for Nissl, myelin, and eight IHC markers.

For specimen 2, the brainstem was dissected at the level of the superior colliculus, where the right and left midbrain were separated at the ventricle during dissection. Hence, the midbrain portion of this specimen was processed separately for histological characterization, with 61 Nissl stained sections along with thalamus (see Table 3). For the 3D reconstruction, the stacked images of the midbrain from both hemispheres were registered to MRI for appropriate alignment. The BFI images were further registered to MRI, resulting in cross-modality registration. We present here IHC datasets from one hemisphere with the corresponding 12 annotated sections (see Table 3).

**Table 3.** The number of manually annotated sections, and the number IHC and myelin stained sections for each specimen.

| Specimen | Age | Nissl | Annotations | Immunohistochemistry (IHC) | Myelin |
| --- | --- | --- | --- | --- | --- |
| 1 | 25 GW | 282 | 78 sections | 39 | - |
| 2 | 9 years | 120 | 91 sections | 530 | 53 |
| 2 | 9 years<br>(Midbrain) | 61 | 12 sections | 72 | 7 |
| 3 | 54 years | 705 | 33 sections | 208 | - |

#### 2.3.1 Immunohistochemical staining

Immunohistochemical staining was performed using the EnVision Flex+ Mouse, High pH (Link) commercial kit (Agilent Dako), designed for detecting primary mouse and rabbit antibodies in formalin-fixed, paraffin-embedded tissue sections.

Prior to immunostaining, sections were immersed in phosphate buffer (PB). Antigen retrieval was achieved by immersing sections in Tris-EDTA buffer and incubating in a water bath at 70°C (fetal brain) or 90°C (adult brain) for 20 min, followed by cooling at room temperature for 20 min. Sections were then washed twice for 5 min each in Tris-buffered saline (TBS).

Endogenous peroxidase was blocked by incubating sections in Peroxidase-Blocking Reagent for 10 min, followed by two 5-min washes in TBS. Nonspecific binding was minimized by incubating sections for 2 h in blocking buffer (0.1 M PB containing 2% bovine serum albumin (ProSpecT), 3% normal goat serum (Vector Laboratories), and 0.25% Triton X-100 (Qualigens).

Sections were incubated for 2 h at room temperature with primary antibodies diluted in blocking buffer: NeuN (Cat. no. ABN78; MilliporeSigma), TH (Cat. no. AB152; MilliporeSigma), OX-A (Cat. no. AB3704; MilliporeSigma), NFH (Cat. no. PA5-34759; Thermofisher), CR (Cat. no. CR7697; Swant), PV (Cat. no. PV27a; Swant), and CB (Cat. no. 2173; CST). This was followed by two 5-min washes in TBS.

Sections were then incubated with HRP-conjugated secondary antibody (EnVision FLEX+ Mouse LINKER) for 30 min, followed by two 5-min washes in TBS. Immunoreactivity was visualized using DAB+ Chromogen in Substrate Buffer. The reaction was terminated with distilled water upon achieving optimal staining intensity.

#### 2.3.2 Myelin staining

Slides were UV-cured prior to staining if required. On Day 1, slides underwent distilled water wash, immersion in pyridine: acetic anhydride for 30–40 min, three washes in 0.5% acetic acid, silver impregnation for 45 min to 1 h, three further 0.5% acetic acid washes (total 10 min), and overnight fixation in 4% paraformaldehyde (2 L). On Day 2, slides received two 0.5% acetic acid washes, immersion in developer. This was followed by 1 min bleaching in 0.2% potassium ferricyanide, 0.5% acetic acid wash, re-immersion in developer until optimal staining. Repetition of bleaching, wash, and immersion in developer was done if needed. Two 5-min 0.5% acetic acid washes, 1 min immersion in 2% sodium thiosulfate, three distilled water washes, dehydration through graded ethanol (50%, 70%, 95%, 100%; 2 min each), 2 min clearing in fresh xylene was done before coverslipping.

### 2.4 Digitization and Quality check

All stained sections were cleared in xylene and coverslipped using an appropriate mounting medium. Digitization was performed using a large-format slide scanner to generate high-resolution whole-section images. The digitized images underwent a quality check procedure which was then utilized for analysis (Verma et al., 2025)

### 2.5 Atlas annotation

All three specimens were manually annotated: 78 sections in the 28 GW specimen, 103 sections in the 9 years old specimen, and 33 sections in the 54-year-old specimen (see Table 3). Additionally, the TH-stained sections have been annotated in the 9 years and 54 years old specimens.

The annotations were made using a custom in-house online annotation tool (Verma et al., 2025), and the brain structures were divided in five general classes: neuronal structures and their subparts, fiber tracts, ventricles and cavities of the brain, transitory structures (e.g., migratory streams; only the 25 GW specimen), and other structures (e.g., ependymal zone). Because of the different angle of cutting of the 25 GW specimen, and the 9 years and 54 years old specimens, respectively (see Table 1), we constructed 2 partially different hierarchically organized nomenclatures. Moreover, the nomenclature constructed for annotation of the 9 and 54 year-old specimens included the catecholaminergic alphanumeric fields (Dahlström & Fuxe, 1964; Manger, 2017; Williams et al., 2022), while the 25 GW specimen includes transitory structures (Bayer & Altman 2004-2006). For both nomenclatures, we also used the human brainstem Paxinos Atlas (Paxinos et al, 2012; 2019; 2023), the human adult brain AIBS Atlas (Ding et al., 2017), and the marmoset Atlas (Paxinos et al., 2012). We list in Appendix Table 1 the parts identified in each specimen.

### 2.6 3D registration

### 2.6.1 Preprocessing

T1-weighted images were used for specimens 1 and 3. For specimen 2, super-resolution reconstruction (SRR) was applied to 2D axial FSE and 2D SAG-FSE acquisitions to generate an isotropic T2-weighted volume (0.8 mm resolution). All datasets were pre-processed using FMRIB Software Library (FSL) 6.0.7 and Advanced Normalization Tools (ANTs) 0.6.2 to ensure a consistent spatial reference. Specifically, the image origin was reset to (0, 0, 0) to standardize coordinate systems, followed by affine header correction and orientation normalization (axis permutation and flipping). For the specimen 1 dataset, a spatial alignment was first performed using anatomical landmarks (the anterior and posterior fontanelles), after which brain extraction was carried out using FMRIB Software Library, Brain Extraction Tool (FSL BET) 6.0.7.22 (fractional intensity threshold = 0.3). For the adult datasets (specimens 2 and 3), spatial initialization used anatomically conserved brainstem landmarks, primarily the pons and medulla oblongata, to create a consistent reference orientation across specimens. Initial alignment involved correcting head orientation and gross rotational offsets to ensure correspondence of the anterior–posterior, superior–inferior and left–right anatomical axes. We visually verified registration quality by checking the correspondence of major neuroanatomical landmarks.

#### 2.6.2 Histological Processing and Multimodal Stacking

Histological sections with the slice thickness (20 µm for specimens 1 and 3; 40 µm for specimen 2) and corresponding BFI were incorporated into the pipeline. Immunohistochemistry images were aligned with their corresponding Nissl-stained sections to form multimodal stacks using Fiji (ImageJ) 2.14.0. Before stacking, each section underwent background removal using adaptive thresholding to eliminate non-tissue regions, followed by histogram matching and adaptive color adjustments to normalize intensity and color variations across the nine distinct tissue stains, thereby removing background artifacts and ensuring both visual and quantitative consistency throughout the imaging stack.

#### 2.6.3 Cross-Modal Registration

Registration across MRI, BFI, and histology was performed using ITK-SNAP v4.4.0 and 3D Slicer v5.8.1. Following a structured two-stage pipeline, all modalities were first brought into a common reference frame. Specifically, the initially independent coordinate systems of the MRI (fixed reference), BFI, and histological sections (moving volumes) were aligned by their spatial orientation and origin. In the second stage, a rigid alignment was performed using manually identified anatomical landmarks (e.g., inferior colliculi and pontine gray). Subsequently, affine registration was applied to account for global differences in scale and shape arising from acquisition and specimen handling. The reconstructed histology and block face volumes were then aligned to the MRI reference space manually, and all datasets were resampled into a common coordinate system. Registration quality was assessed through visual verification of anatomical boundary alignment across modalities.

Initial spatial relationships are established using down sampled 16-µm resolution images, leveraging SIFT (Scale-Invariant Feature Transform) to extract local features for multi-modal alignment. Rigid registration parameters calculated sequentially relative to a designated seed section ($I_0$) serving as the global anatomical anchor are propagated across the volume to yield a cohesive 3D structure. Applying these transforms to the full dataset generates a 60-µm isotropic histology stack, ensuring the structural continuity required for integration with 3D MRI and BFI coordinates. Within the visualization interface, a strict pairing mechanism maintains pixel-level correspondence among adjacent Nissl, IHC and myelin-stained sections. This precise co-registration enables co-localized analysis of cytoarchitecture, protein expression, and fiber pathways across the 9-year-old specimen dataset.

#### 2.6.3 Multimodal Data Registration and Visualization

The multimodal neuroimaging brainstem pipeline integrates macroscale volumetric data with microscale cellular details through a rigorous co-registration framework. The process begins with MRI to establish the global anatomical reference, followed by BFI, which captures the physical geometry of the tissue block during the sectioning process. These BFI images are crucial because they lack the non-linear distortions typically found in thin histology sections, acting as a spatial "bridge" between the 3D MRI volume and 2D microscopy.

### 2.7 Web-Based Low-Footprint Viewer

By establishing precise spatial correspondence across these modalities, the viewer enables a seamless transition from gross brain structures in the MRI to features at the cellular level. This unified coordinate system allows navigation across adjacent histological sections, from Nissl to the seven IHCs, for all the annotated sections simultaneously. Figure 2 shows the online viewer for 9 years old specimen with more than 700 brainstem sections.

**Figure 2.**
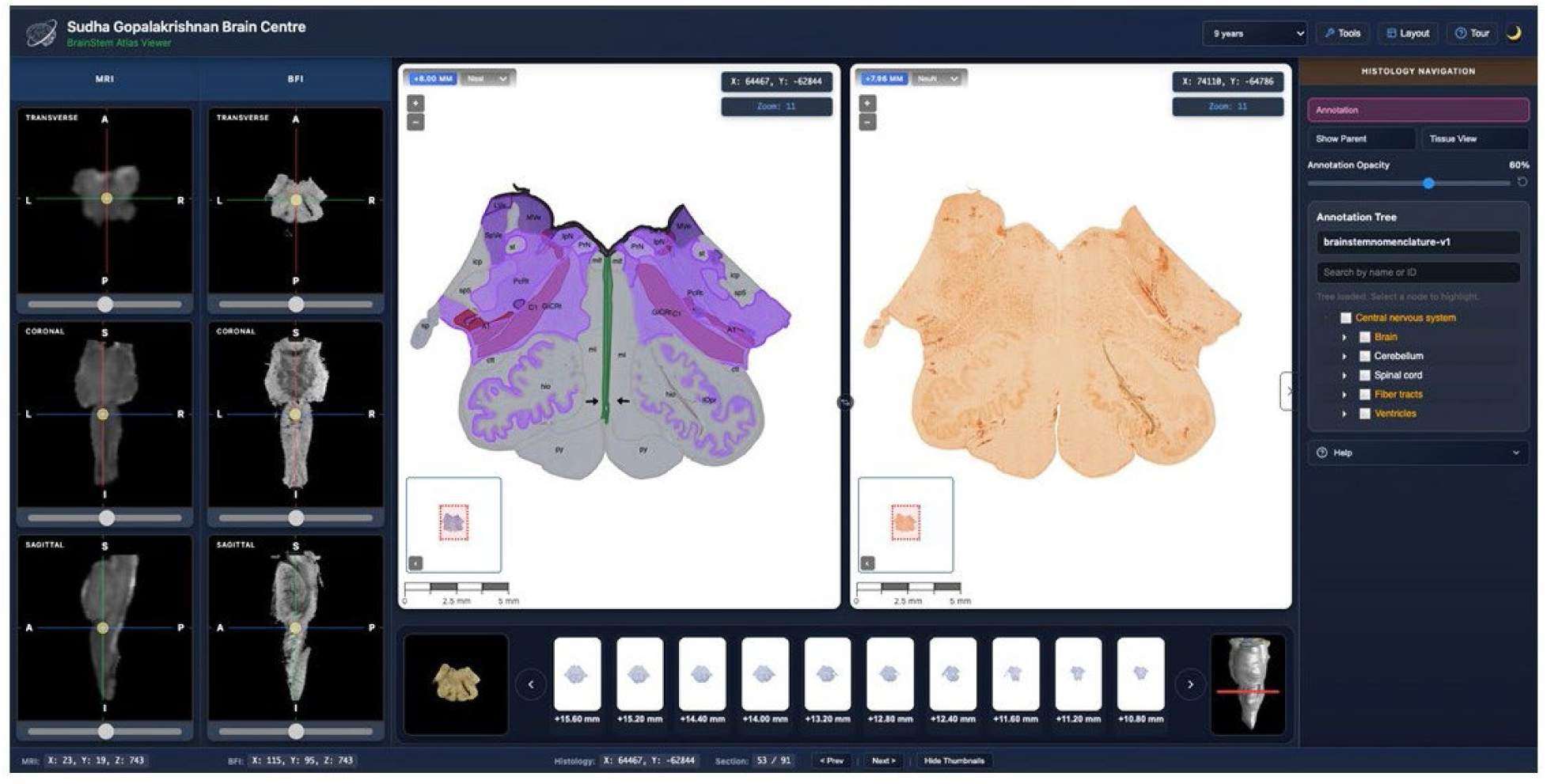
The ANCHOR online viewer seamlessly integrates multimodal neuroanatomical information across different modalities and scales.

Individual section images exceed tens of gigabytes, requiring substantial computation to reconstruct the multi-hundred-section volume. Raw 8-bit RGB TIFF scans were loaded as pixmaps, transformed, color-adjusted, and converted into multi-resolution image pyramids using the *libvips* dzsave command. This tiling strategy minimizes data overhead by serving only the specific resolutions requested by the web viewer. Furthermore, the viewer dynamically maps the spatial correspondence between MRI, BFI and various histological stains.

For the 9 years old specimen, the online viewer offers synchronized, side-by-side panes, that align with identical spatial coordinates, enabling users to concurrently zoom, pan, and examine high-resolution cellular insets of Nissl and IHC stains while preserving macroscopic context through co-registered MRI and BFI. This multi-stain correlation is extremely beneficial, because it correlates Nissl distinct feature with the IHC expression patterns, and with myelin fiber pathways. Cross-referencing these complementary visual cues enables researchers to confidently identify uncertain anatomical borders of complex neuronal parts and validate macro-to-micro registration across imaging modalities.

### 2.8 3D Visualization

The histology 3D reconstructions, which are calculated at 16-µm in-plane resolution and 20-µm slice separation, are saved in Nifti format as a 3D RGB volume in the same coordinate space as the MRI. After inverting the contrast and creating a grayscale volume with a 4-µm in-plane resolution and a 20-µm slice separation, the raw array data was stored. The raw volume is represented using Nvidia IndeX software, which includes direct volume rendering, shaded depiction of orthogonal planes, and pseudo colouring with a colour map theme that represents cell density from blue (low), to orange (high; see videos in the Supplementary Information).

## 3. Results

Overall, we have identified about 280 parts (nuclei, fiber tracts and ventricles) in all three specimens. In the 25 GW specimen we have annotated 189 parts, in the 9 years old specimen 208 parts, and in the 54 years specimen 182 parts (see Appendix Table 1). Table 3 summarizes the number of sections per specimen. Table 4 summarizes the number of parts identified in each specimen.

**Table 4.**
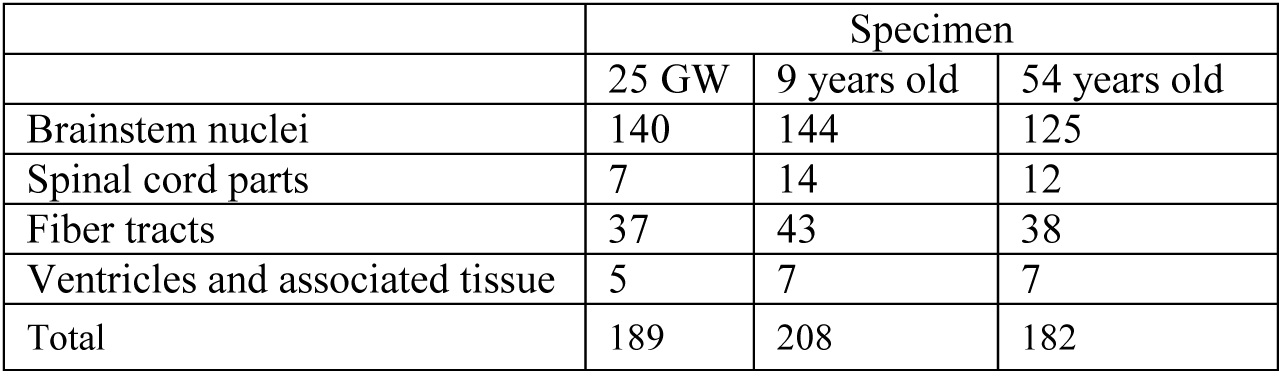
The number of annotated parts per specimen and type of part.

A detailed description of the cytoarchitecture and chemoarchitecture of each specimen is beyond the scope of this article. Therefore, we briefly describe the cytoarchitecture and IHC staining patterns in all three specimens. We also briefly describe the cytoarchitecture of the pretectum in the 25 GW specimen, because the available literature on this region is scarce (Verma et al., 2025).

### 3.1 NeuN

NeuN labeled the brainstem nuclei consistently in all three specimens. Examples are shown in the spinal cord (SP), where its laminae are labeled in all three specimens. (see Figs. 3-5).

**Figure 3.**
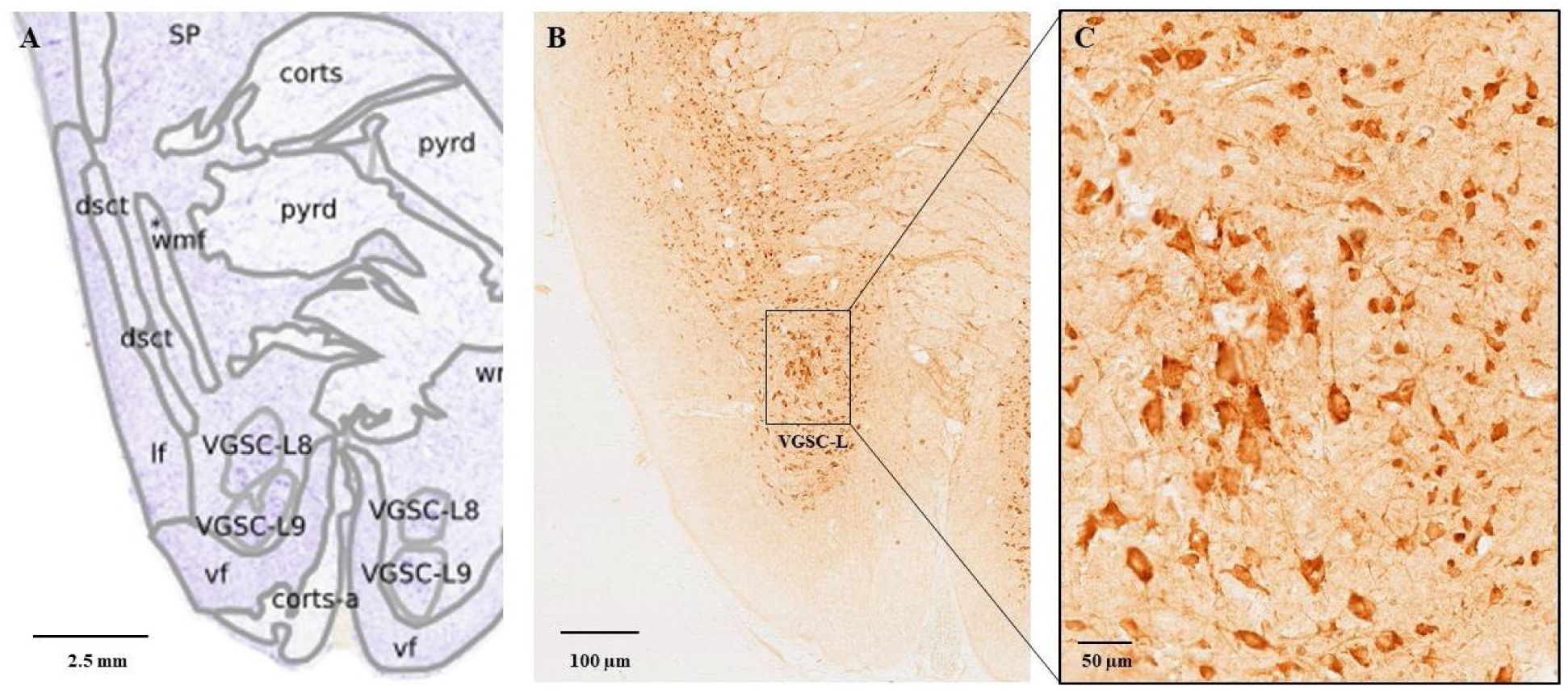
The NeuN label in the layers 8-9 of the SP, 25 GW specimen. A. Nissl stained and annotated section at the level of SC. B. The corresponding NeuN stained section. C. Details of the labeled neurons in layers 8-9 of SP.

**Figure 4.**
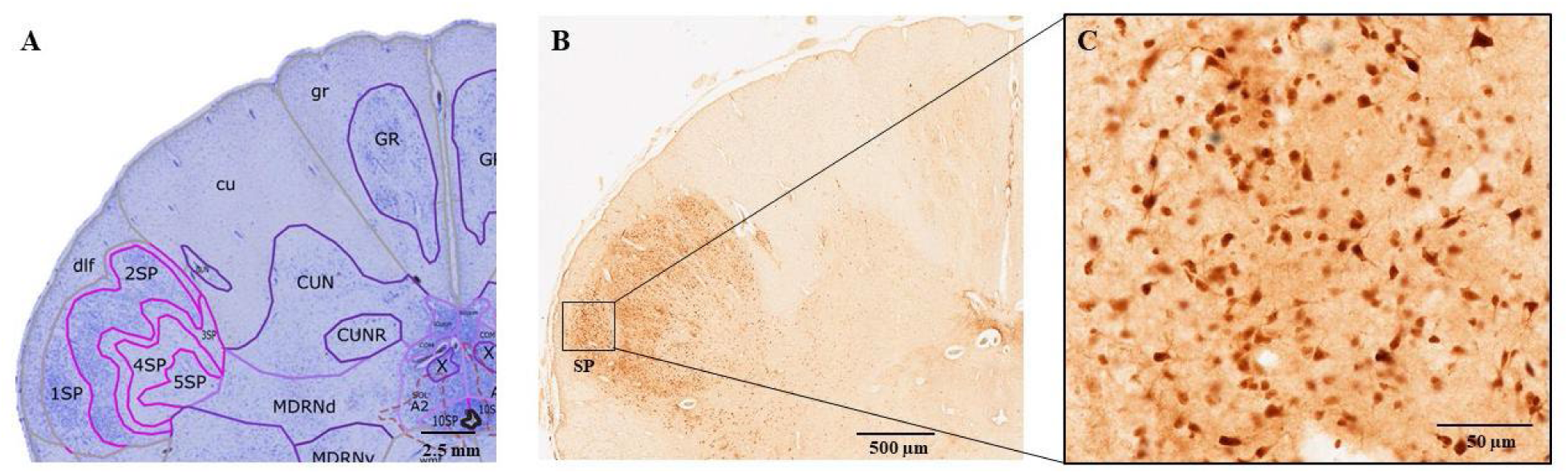
NeuN label in layers 2-4 of SP in the 9 years old specimen. A. Nissl stained and annotated section (obex: - 11.2 mm). B. The corresponding NeuN stained section (obex: −11.48 mm). C. Details of the labeled neurons in layers 2-4 of the SP.

**Figure 5.**
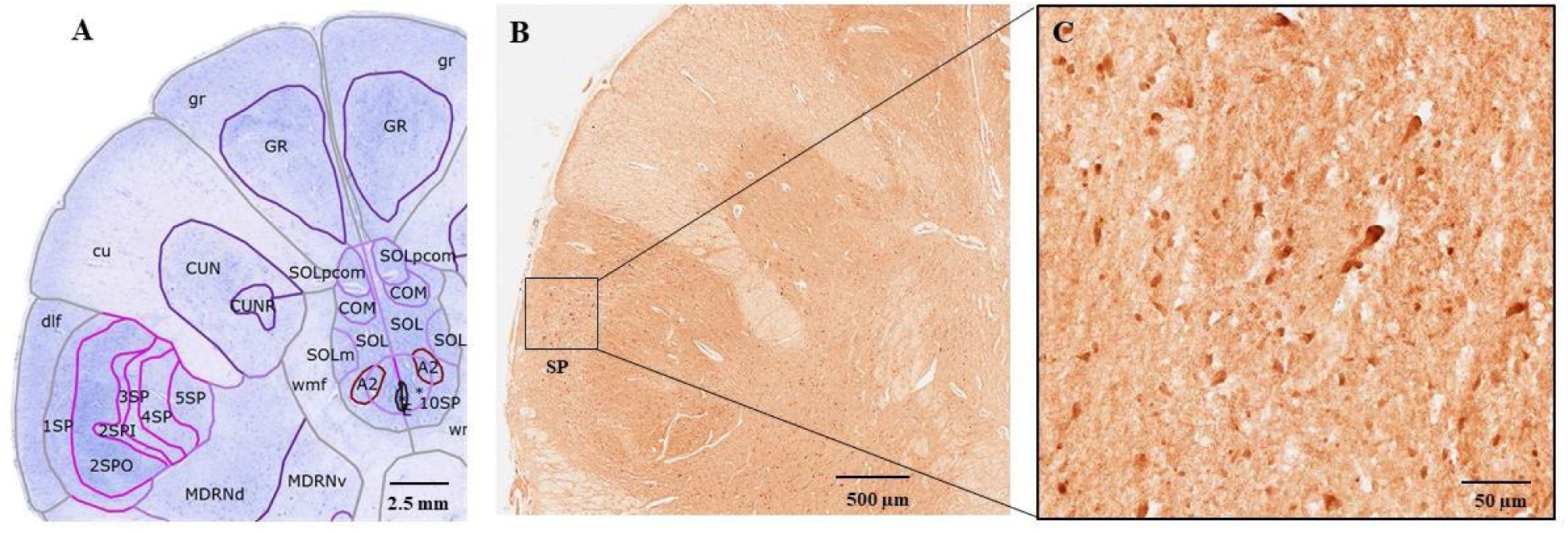
NeuN label in layer 2 of the SP in the 54 years old specimen. A. Nissl stained and annotated section (obex: −7.7 mm). B. The corresponding NeuN stained section (obex: −6.74 mm). C. Details of the labeled neurons in layer 2 of the SP.

### 3.2 Tyrosine hydroxylase

The TH staining patterns were broadly similar in all three specimens. In the medulla, the A1, A2, C1 and C2 fields were identified, as well as area postrema (AP). These fields include labeled neurons and fibers. In the pons, the locus coeruleus nuclear complex was comprised of several subdivisions. The smallest of these was the fifth arcuate nucleus (A5) (Williams et al., 2022), that was found rostrolateral to the facial nerve nucleus. The neurons of the locus coeruleus (LC, A6) were intensely stained, with the neurons exhibiting a diagonal orientation on a latero-medial axis (Figs. 6-8).

**Figure 6.**
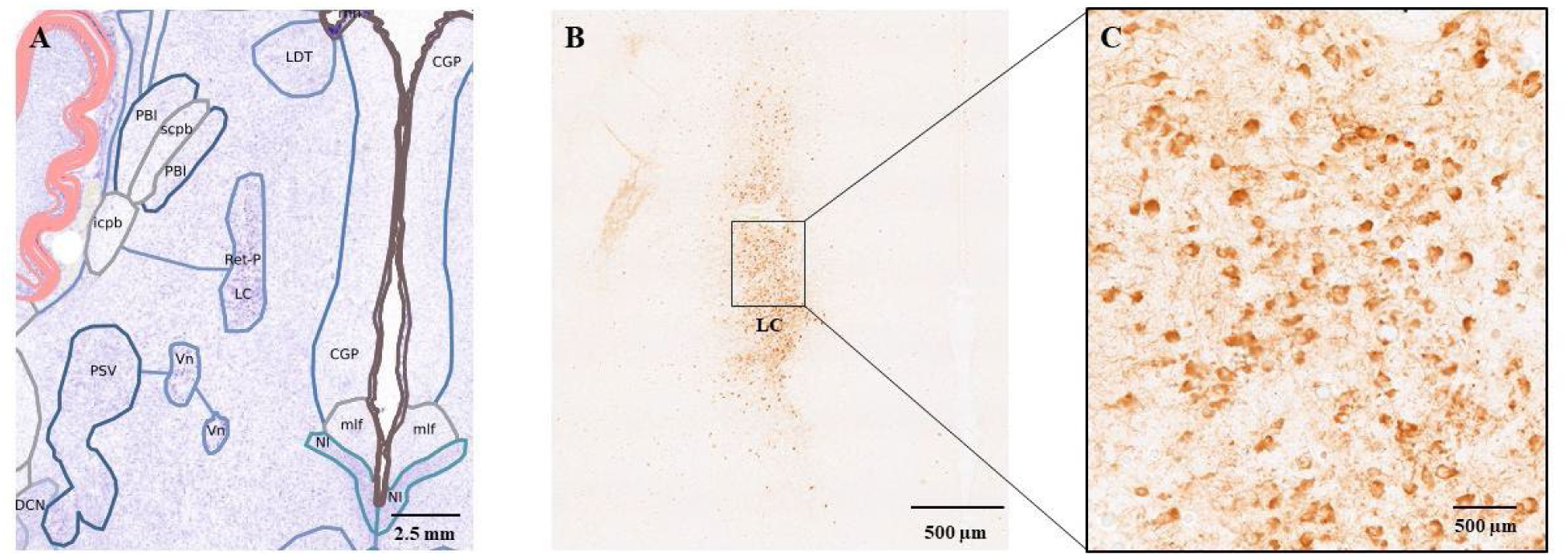
The A6 and A7 fields in the 25 GW fetal brain. A. Nissl stained and annotated section. B. The corresponding TH-labeled section. The A6 and A7 fields are continuous and have a superior to inferior orientation. C. Details of TH-labeled neurons, that tend to be round and moderately intensely stained.

**Figure 7.**
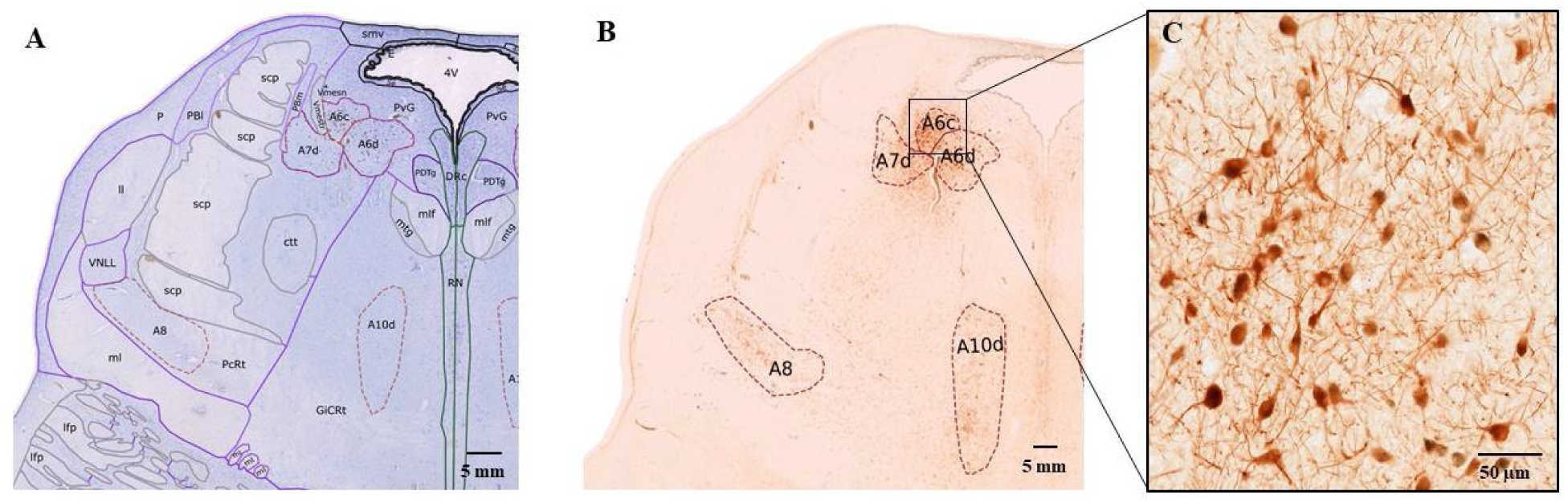
The A6 and A7 fields in the 9 years old specimen. A. Nissl stained and annotated section at the level of pons (obex: +30.4 mm). B. The corresponding TH-labeled section, which includes A6c, A6d, A7d, A8, and A10d TH fields (obex: +30.32 mm). C. Detail of A6c, where TH-labeled neurons are round-to-oval, intensely stained, and oriented dorsomedial to ventrolateral.

**Figure 8.**
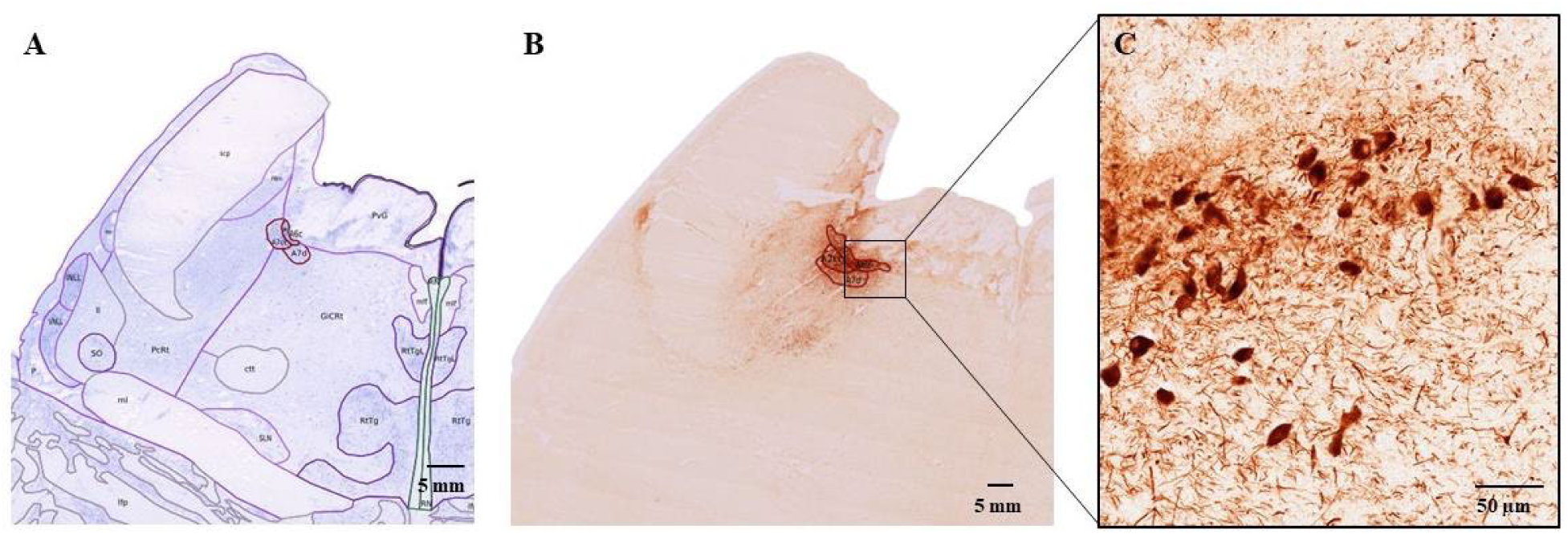
The A6 and A7 fields in the 54 years old specimen. A. Nissl stained and annotated section at the level of A6 and A7 fields (obex: +20.7 mm). B. The corresponding TH-labeled section, which includes A6c, A6d, and A7d fields (obex: +21.56 mm). C. Labeled neurons in the A6 and A7 field, which tend to be round-to oval, intensely stained, and oriented dorsomedial to ventrolateral.

Based on the orientation and density of the labeled neurons, the TH LC field, located entirely within the periventricular gray matter, can be further subdivided in a compact part (A6c), and a diffuse part (A6d; Williams et al., 2022). Ventrolateral to LC, separated by the passage of the fifth mesencephalic tract and within the pontine tegmentum, we identified the locus subcoeruleus (A7), with two subdivisions: a compact (A7sc), and a diffuse part (A7d; Figs 6-8). The cell bodies of the labeled neurons in A7d tend to be round (Figs. 6-8). TH-labeled descending fibers and originating in the A7 fields were also observed (Figs 6-8). In the 25 GW specimen, the A6 and A7 fields form a continuous structure, not being separated by the passage of the fifth mesencephalic tract, oriented along a superior-inferior axis (Figure 6). The soma of the neurons are round and do not have the specific dendritic orientations observed in the adult specimens.

In all three specimens labeled fibers originate from the A6 and A7 fields, curve around the superior cerebellar peduncle (scp), and end in parabrachial nucleus, lateral part (PBl), and very close to the scp (Figs.9-11).

**Figure 9.**
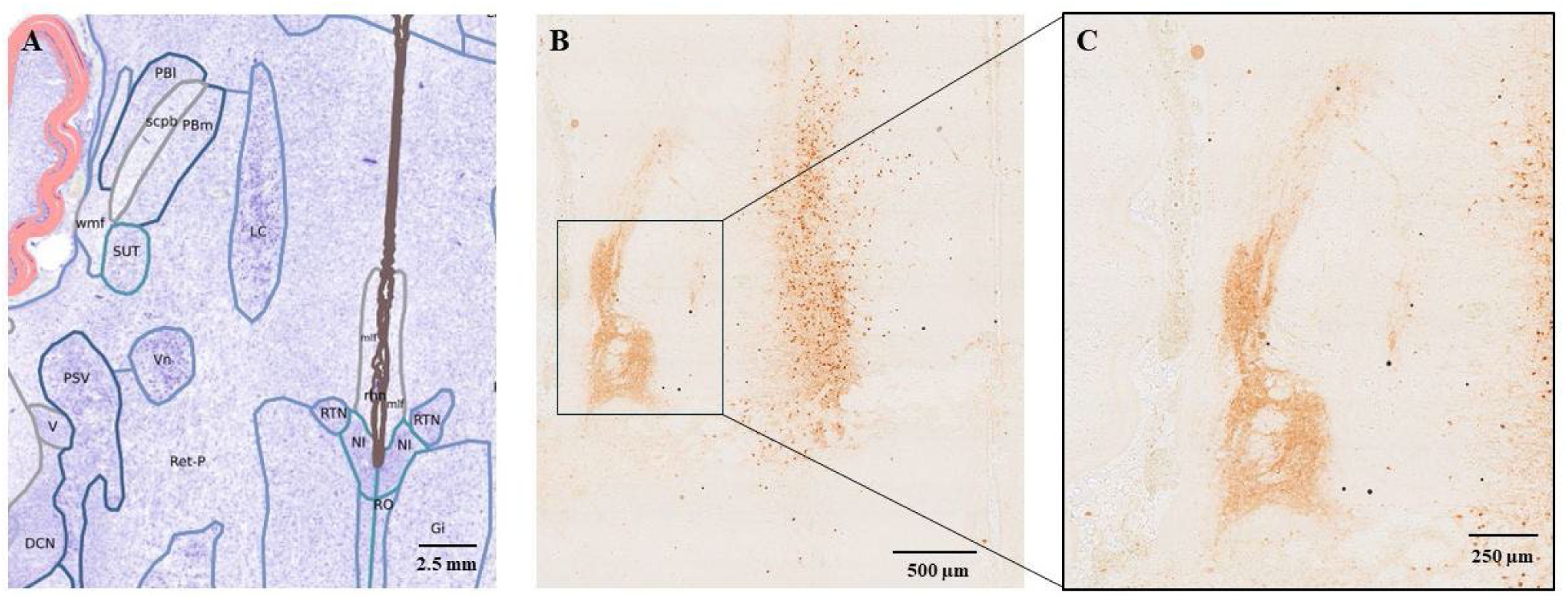
Ascending fibers from the A6 and A7 fields observed in the PBl, 25 GW specimen. A. Nissl stained and annotated section at the level of pons. B. The corresponding TH stained section. C. Terminal field in the PBl, adjacent to the scp.

**Figure 10.**
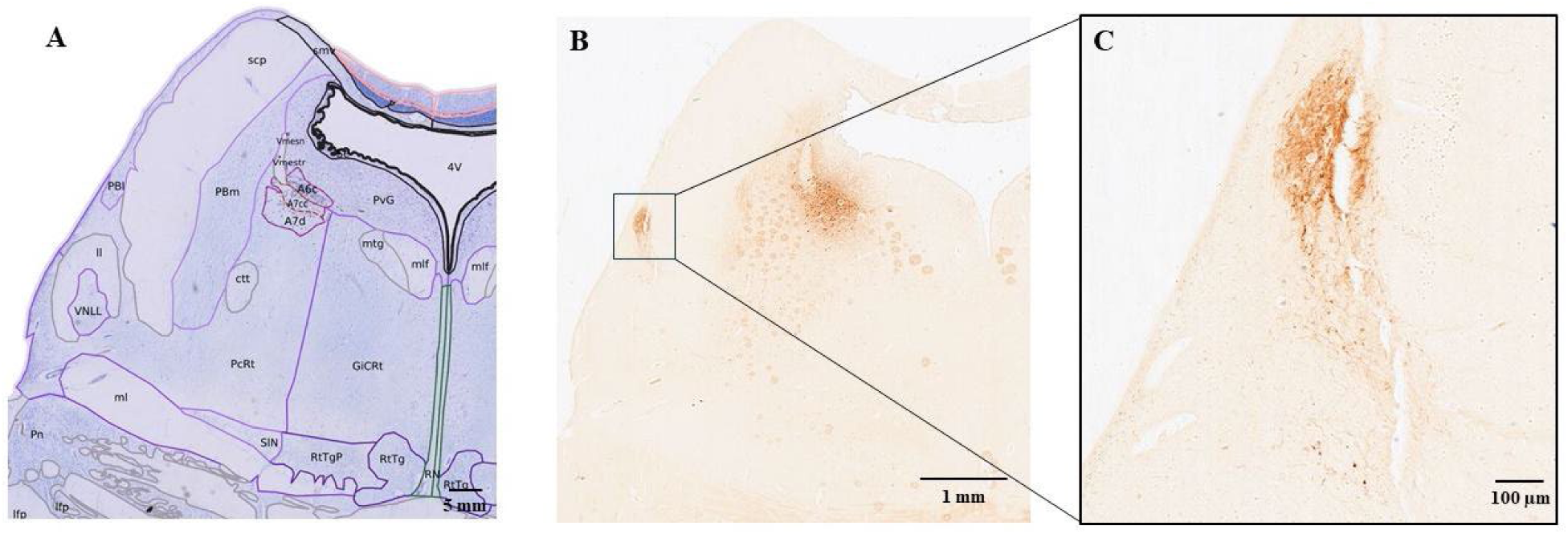
Ascending fibers from the A6 and A7 fields observed in the PBl, 9 years old specimen. A. Nissl stained and annotated section (obex: +26.80 mm). B. The corresponding TH stained section (obex: +26.72 mm). C. Detailed view of the terminal field in the PBl, adjacent to the scp.

**Figure 11.**
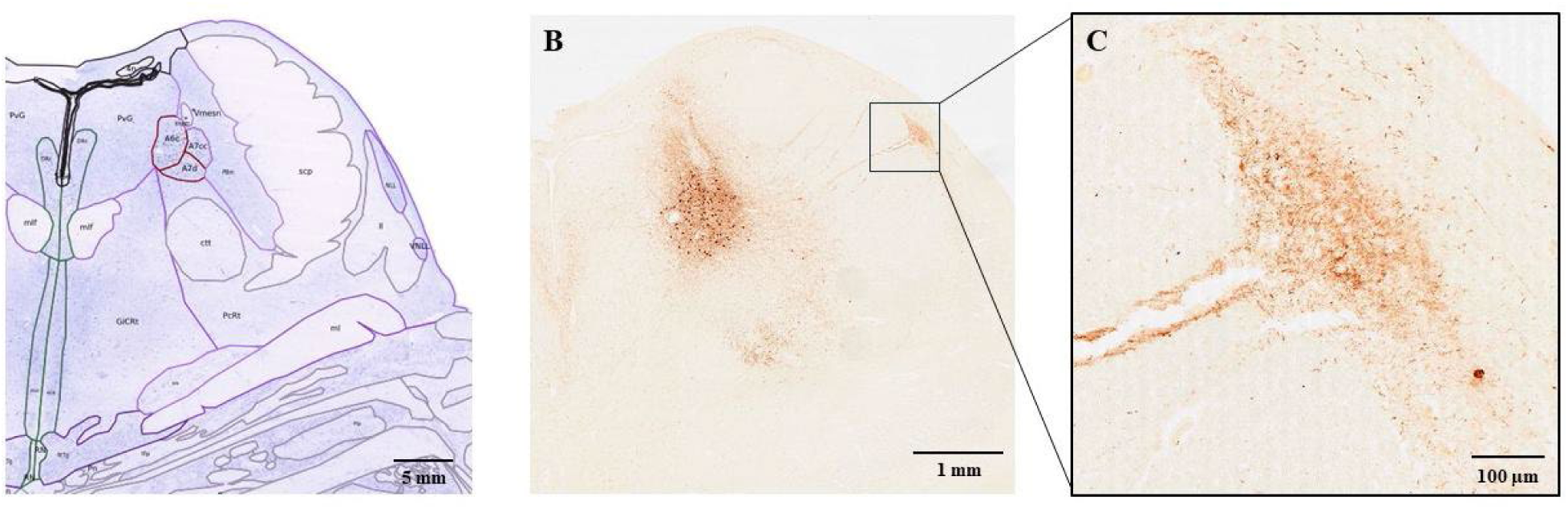
Ascending fibers from the A6 and A7 fields observed in the PBl, 54 years old specimen. A. Nissl stained and annotated section (obex: +30.40 mm). B. The corresponding TH-stained section (obex: +30.32 mm). C. Detailed view of the terminal field in the PBl, adjacent to the scp.

Furthermore, in the 54 years old specimen TH labeled fibers were observed in the scp, with an orientation orthogonal to that of the scp fibers (see Fig. 11).

In the midbrain, we identified the A8 field (retrorubral area), located caudal and lateral from the red nucleus (R). At its largest part the A8 is bounded medially by the medial lemniscus, and it is bordered laterally by the superior cerebellar peduncle decussation (xcsp; see Fig. 12).

**Figure 12.**
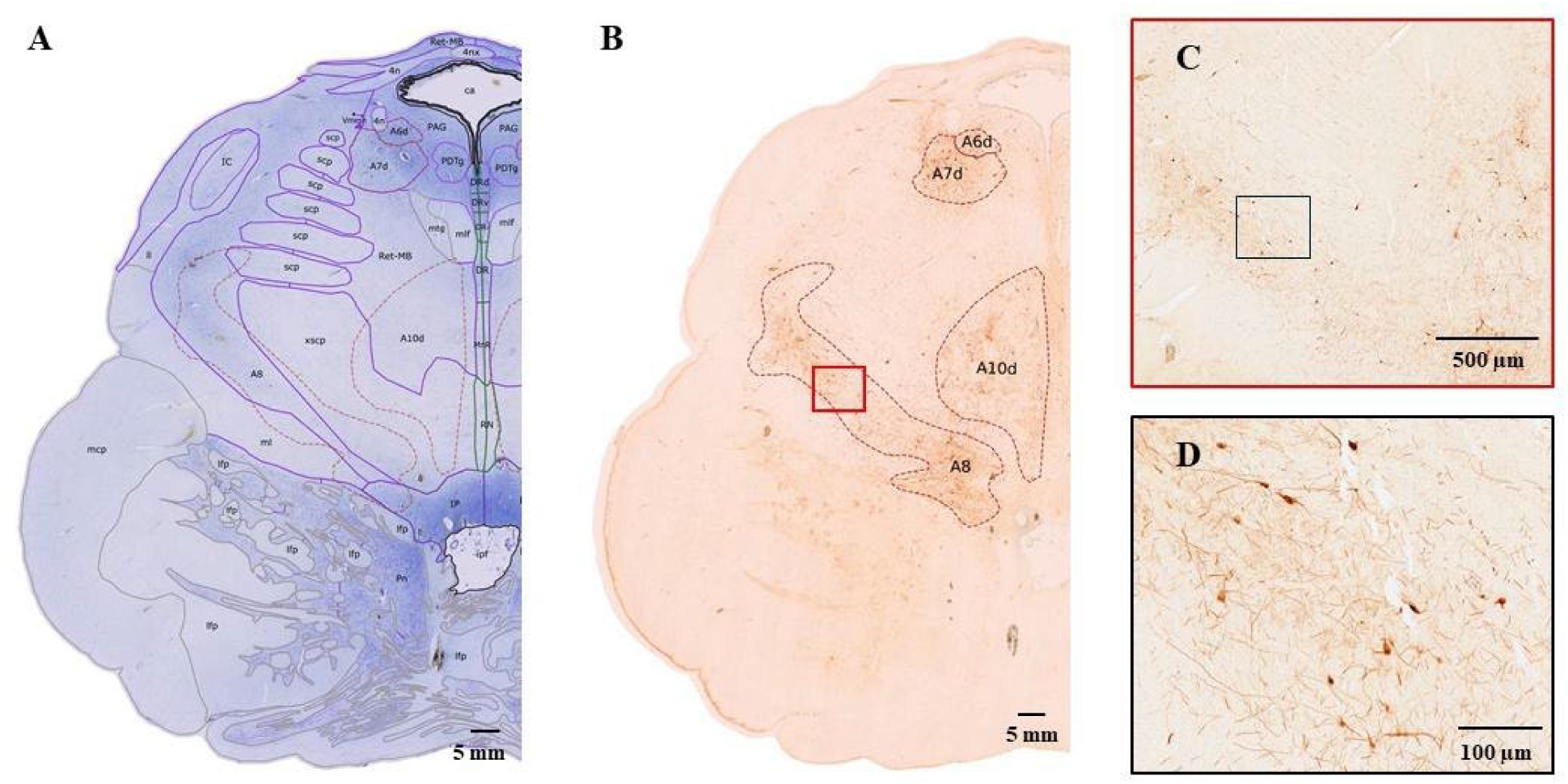
The retrorubral area in the 9 years old specimen. A. Nissl labeled and annotated section (obex: +32.80 mm). B. The closest TH-labeled section (obex: +32.72 mm). C, D. Details of the labeled neurons and fibers in the retrorubral field.

A8 as identified here is more extensive than previously described (Halliday et al., 2012). The substantia nigra included a large TH-labeled field subdivided in pars compacta (A9pc), lateral (A9l), medial (A9m) and ventral (A9d) subparts. In the midbrain, the ventral tegmental area (VTA) could be subdivided in dorsal (A10v), and dorsocaudal (A10dc) parts (Williams et al., 2022). TH-positive neurons also have been identified in the parabrachial pigmented nucleus in adult specimens. Finally, TH-labeled fibers have been observed entering the red nucleus in the 9 and 54 year old specimens.

### 3.3 Calbindin

In the cervical spinal cord, CB staining labeled intensely layers 1-3. (see Figs. 13-15).

**Figure 13.**
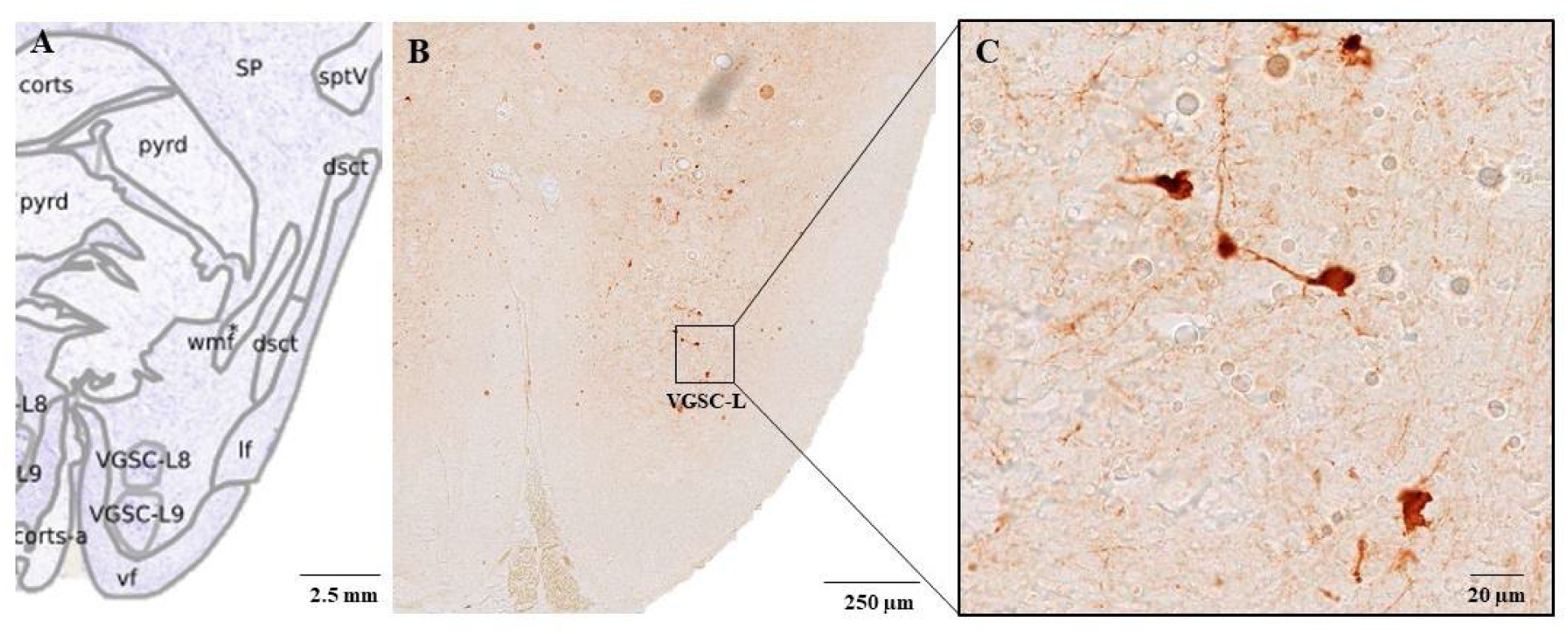
CB labeled neurons in the layers 8-9 of SP, in the 25 GW specimen. A. Nissl labeled and annotated section. B. The corresponding CB-labeled section. C. Details of labeled neurons and fibers in the layers 8-9 of SP.

**Figure 14.**
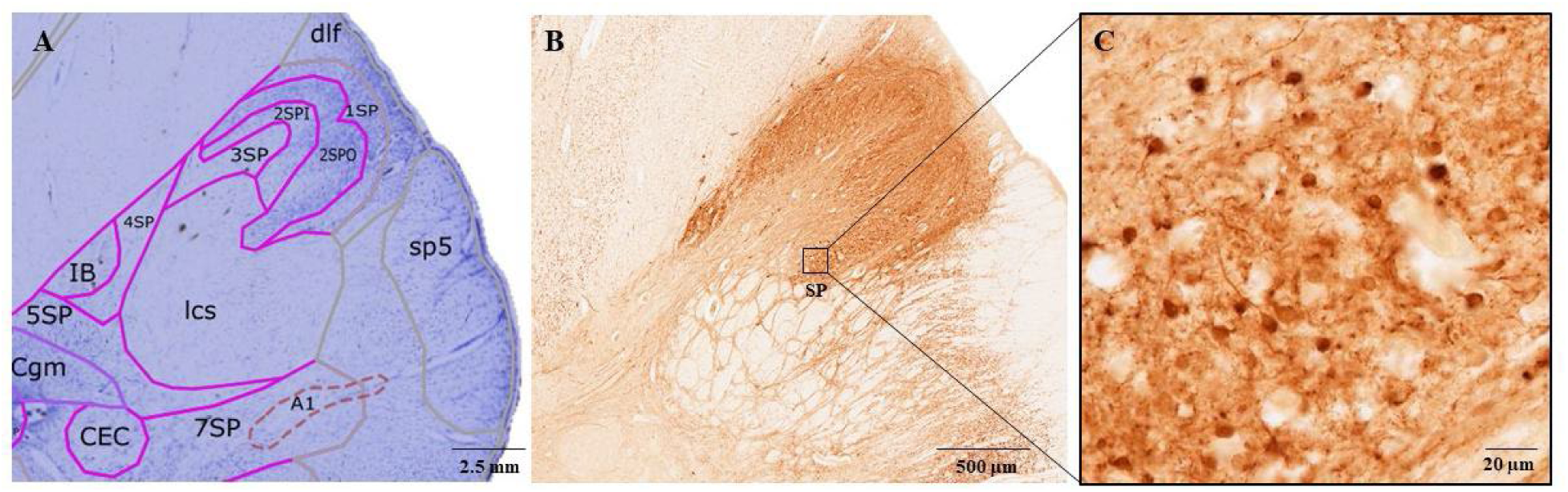
CB labeled neurons and neuropil in layers 2-3 of SP, in the 9 years old specimen. A. Nissl labeled and annotated section (obex: −11.20 mm). B. The corresponding CB-labeled section (obex: −11.42 mm). C. Details of neurons, fibers, and neuropil in layers 2-3 of SP. Compare with Figure 13C.

**Figure 15.**
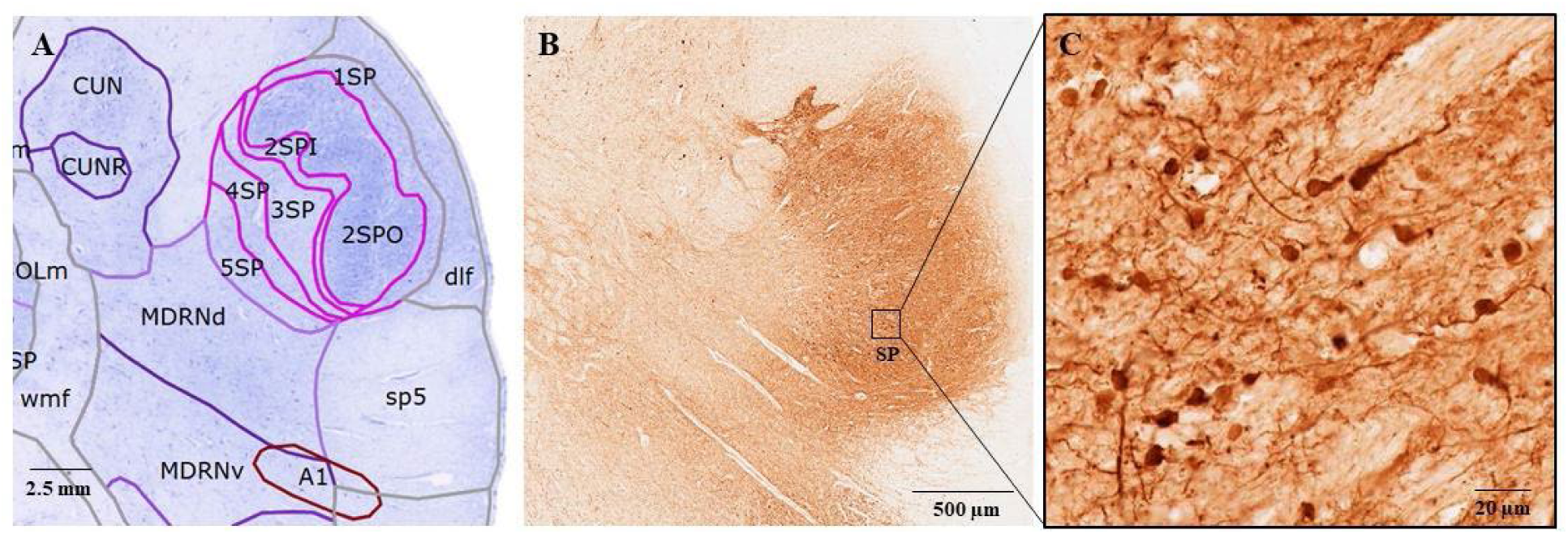
CB labeled neurons and neuropil in layers 2-3 of SP, in the 54 years old specimen. A. Nissl labeled and annotated section (obex: −7.68 mm). B. The corresponding CB-labeled section (obex: −6.74 mm). C. Details of neurons, fibers, and neuropil in layers 2-3 of SP. Compare with Figure 13C.

In the medulla, the gracile nucleus (GR) was labeled, especially anteriorly. Labeled neurons and neuropil were present in the vestibular and spinal nuclei. In the medulla, the inferior olive (IO) was labeled in all three specimens (see Fig. 16). Small neurons with round soma could be identified in the main nucleus of the IO, as well as fiber tracts that presumably start in the hilus of the IO, cross the spinal nucleus of the trigeminal nerve (SP5), end in the inferior cerebellar peduncle (icp), and which are collectively described as the olivo-cerebellar fibers (Parent, 1996; see Fig. 16).

**Figure 16.**
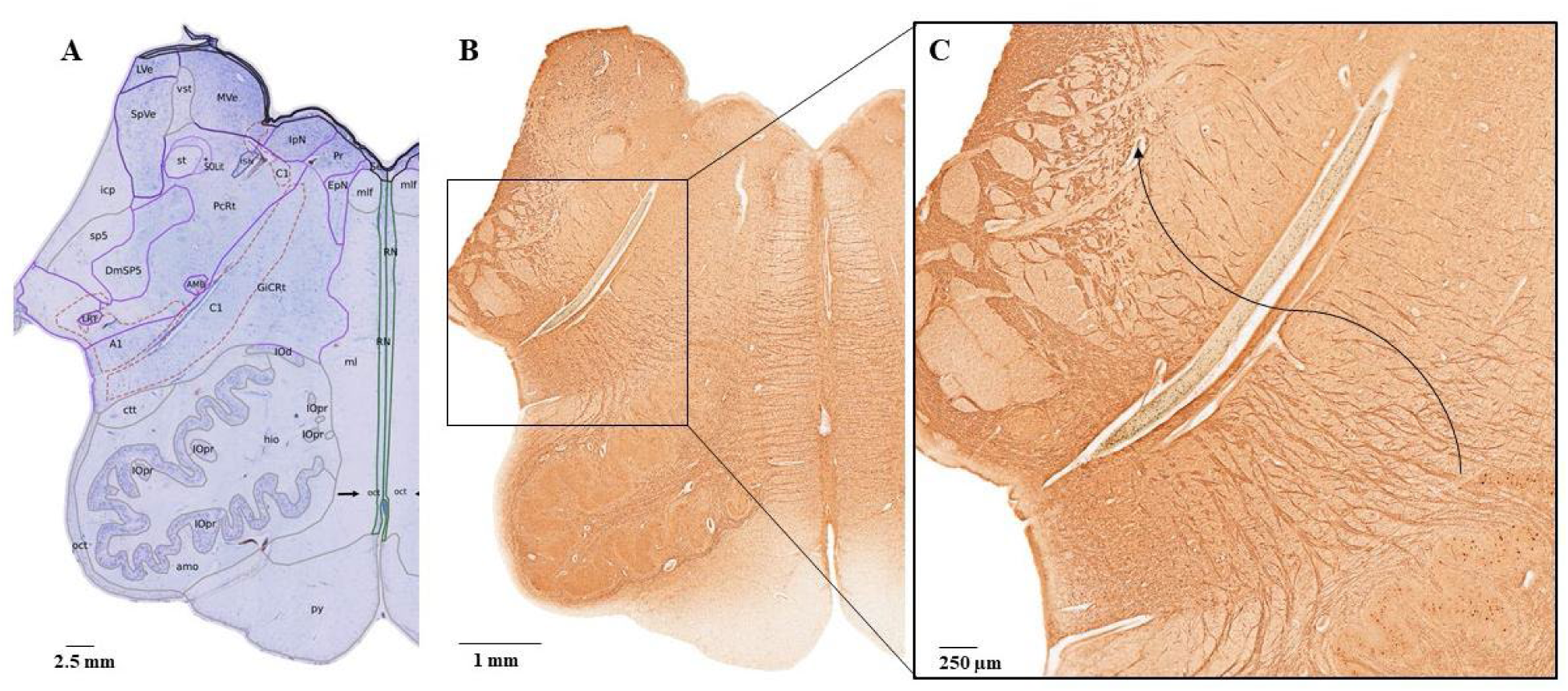
CB-labeled fibers at the level of IO in the 9 years old specimen. A. Nissl stained and annotated section at the level IO (obex: +8.0 mm). B. The corresponding CB-stained section (obex: +7.72 mm). Longitudinal CB-labeled fibers extend from IO to the midline. C. Detailed view of ascending CB-stained fibers, indicated by the arrow, that start from IO, cross SP5 and end in the icp.

Longitudinal CB-immunoreactive fibers have also been observed at the IO levels (see Fig. 16B). In the pons, the SO was labeled in the 25 GW and 9 years old specimens. An intense field of labeled neuropil and few neurons was observed above the genu of the facial nerve (7n). In the 25 GW specimen, the dorsal raphe (DR), and the dorsal tegmental nucleus (DTN) included labeled neurons and fibers. The medial longitudinal fasciculus (mlf) included CB labeled fibers. In the midbrain, the inferior colliculus (IC) was devoid of label.

### 3.4 Calretinin

The CR immunostaining patterns were similar with those produced by CB staining. In the SC, layers 1 and 2, and more anteriorly layer 5, were intensely labeled. In the medulla, moderately stained fields were identified in GR and in the cuneate nucleus (CU). The arcuate nucleus of the medulla (AR) was intensely labeled, and stained neuropil was identified in the posterior solitary nucleus (SOL). The IO was moderately labeled, and fiber tracts originating from the hilus of IO to the icp were also labeled (see Fig. 17).

**Figure 17.**
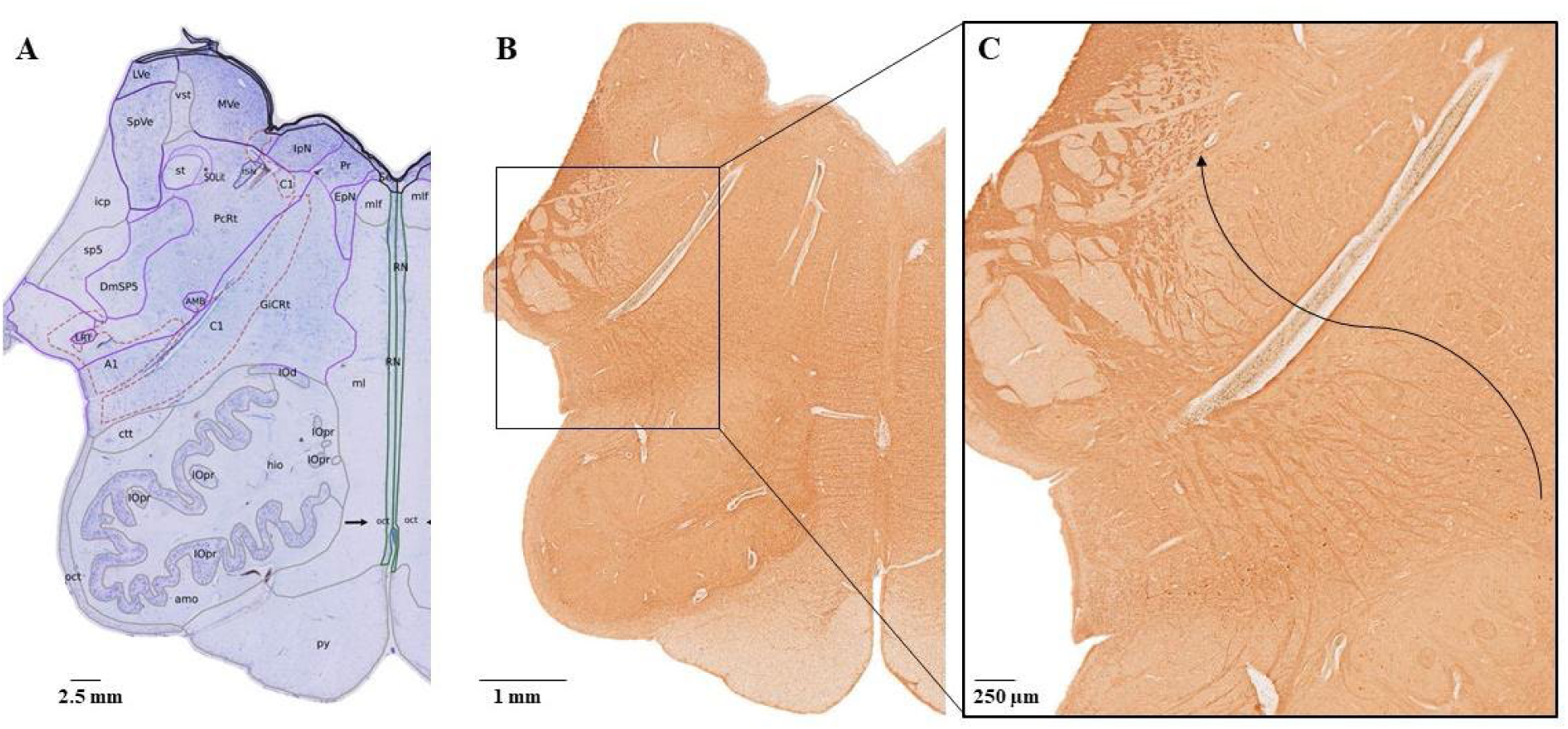
CR-labeled fibers at the level of IO in the 9 years old specimen. A. Nissl stained and annotated section at the level IO (obex: +8.0 mm). B. The corresponding CR-stained section (obex: +7.8 mm). Longitudinal CR labeled fibers extend from IO to the midline. C. Detailed view of ascending CR-stained fibers, indicated by the arrow, that start from IO, cross SP5 and end in the icp.

In the pons, the SO was labeled in a similar fashion with CB staining. The ll was labeled as a fan-like set of fibers exiting the SO, and traveling laterally, and dorsally. The CB+ field above the genu of the 7^th^ nerve was also present in CR staining. Finally, in the midbrain the IC was devoid of label.

### 3.5 Parvalbumin

In the cervical spinal cord, PV+ immunostaining was identified in lamina 1 and 5. In the medulla, GR and CU included dense PV fields. In the 9 years and 54 years old specimens, the AR was labeled. In SOL, the immunostaining was restricted to a few nuclei. The IO was labeled, especially its dorsal part (IOd). In the dorsoposterior part of the pons, the vestibular nuclei were intensely labeled. Also in pons, the facial nucleus was intensely labeled, and a small field of labeled neurons were identified in its dorsal part. An intensely stained region of labeled neurons and neuropil was visible in SO (see Figs. 18-20).

**Figure 18.**
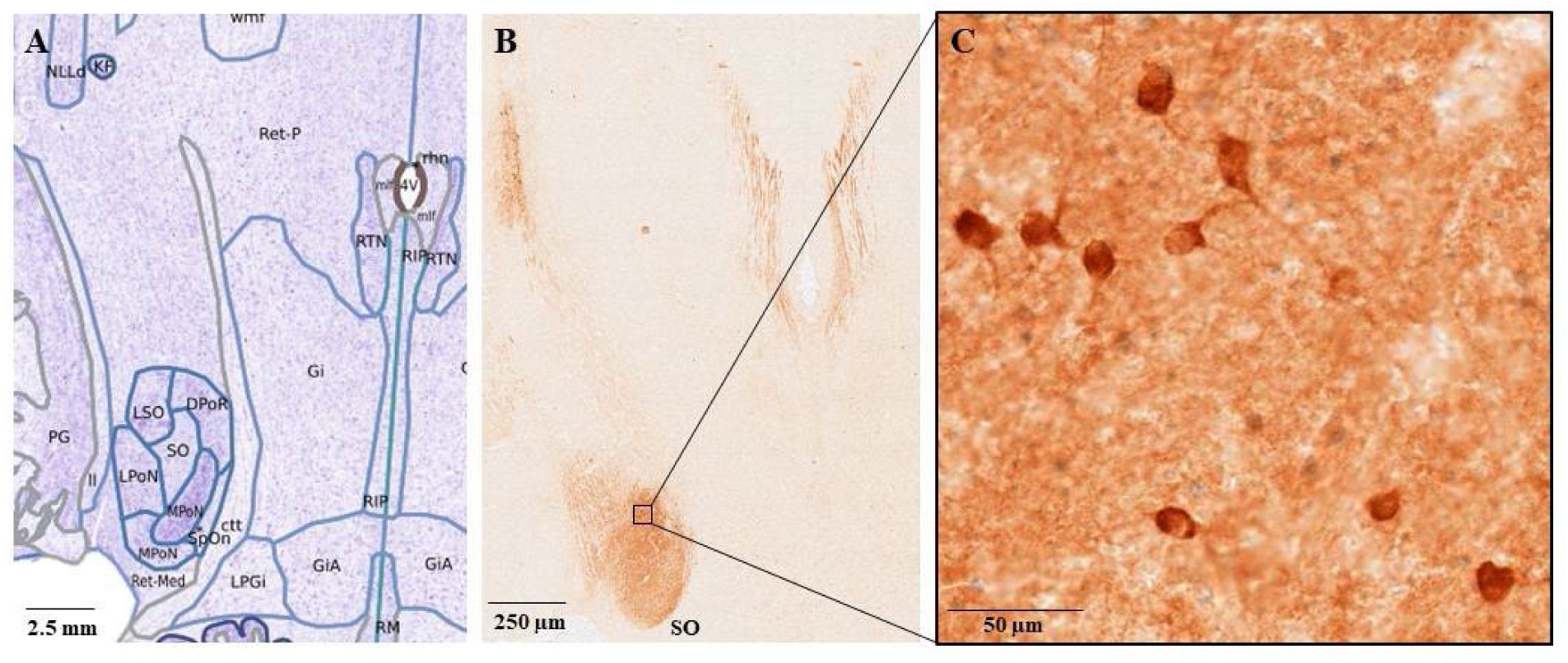
PV labeled neurons and neuropil in the SO, 25 GW specimen. A. Nissl stained and annotated section at the level of SO. B. The corresponding PV-stained section. C. Details of labeled neurons and neuropil in SO.

**Figure 19.**
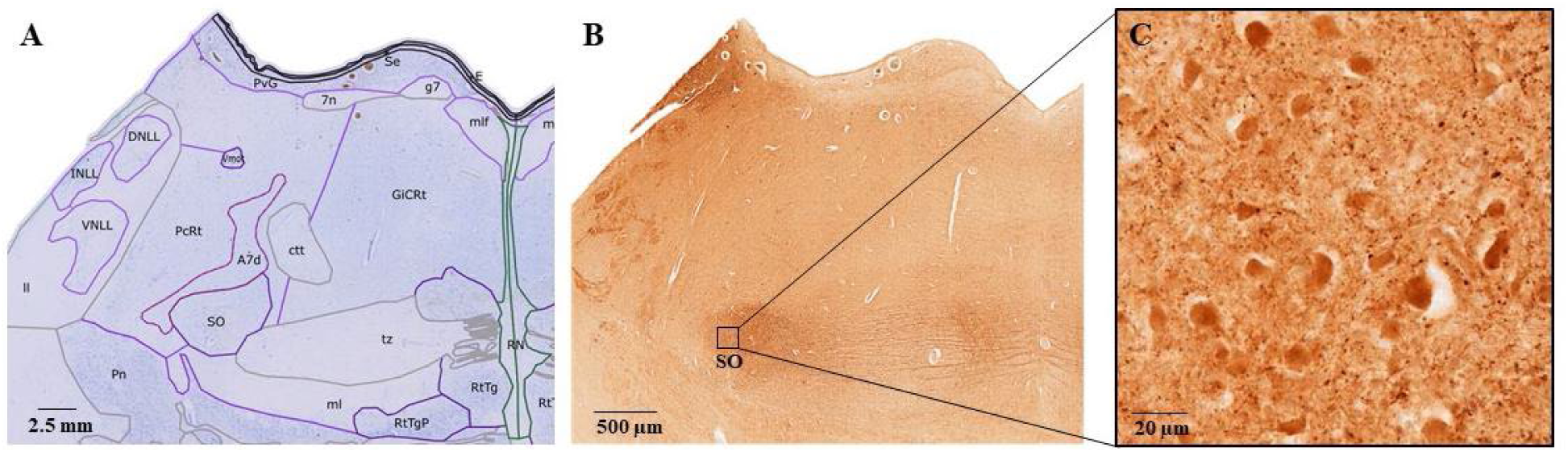
PV-labeled neurons and neuropil in the SO, 9 years old specimen. A. Nissl stained and annotated section at the level of SO (obex: +18.4 mm). B. The corresponding PV-stained section (obex: +18.16 mm). Fibers leaving the SO and traveling towards the midline through the trapezoid body, presumably to the raphe nuclei The NLL also include PV-stained neurons and neuropil. C. Details of labeled neurons and neuropil in SO.

**Figure 20.**
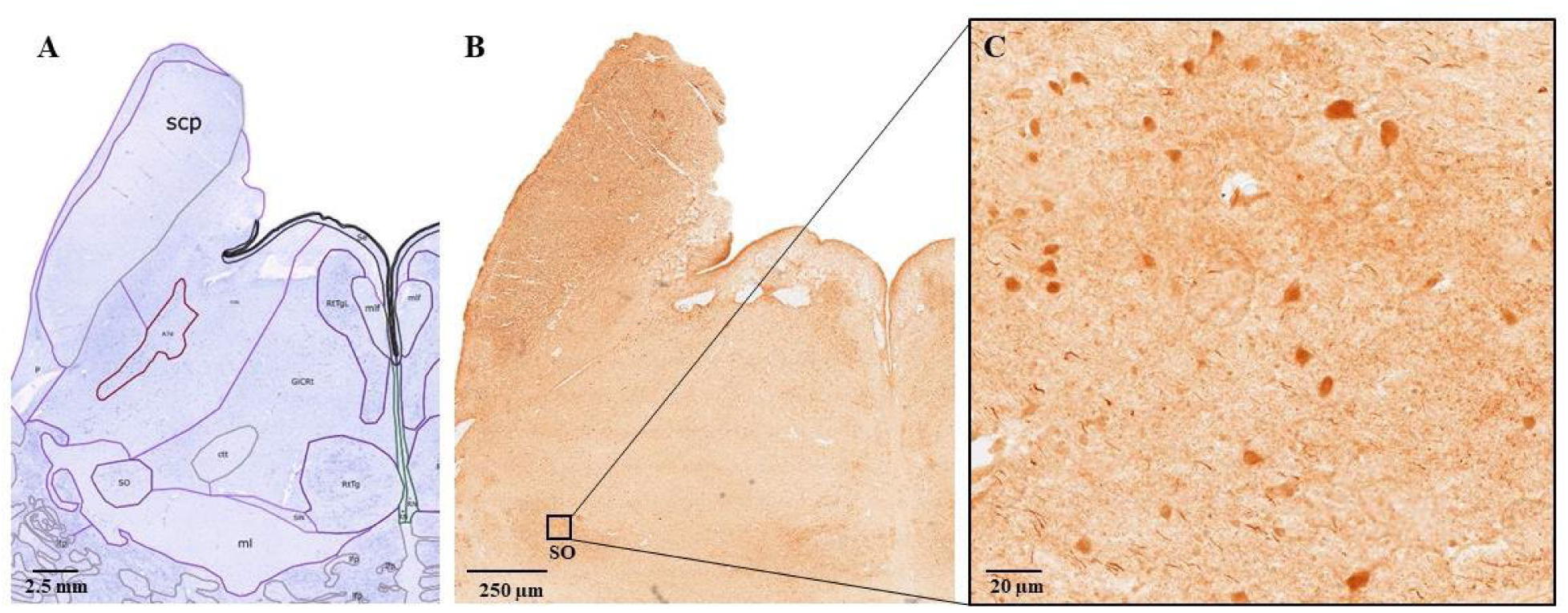
PV-labeled neurons and neuropil in the SO, 54 years old specimen. A. Nissl stained and annotated section at the level of SO (obex: +17.3 mm). B. The corresponding PV stained section (obex: +17.60 mm). C. Details of labeled neurons and neuropil in SO.

Labeled longitudinal fibers were identified between the SO and midline, traveling through the trapezoid body and presumably the raphe nuclei, where small neurons and neuropil were present (see Fig. 19). The nuclei of the lateral lemniscus (NLL; see Fig. 19) included labeled neurons and neuropil. Labeled fibers could be seen in the lateral and medial lemnisci. In the adult specimens, small interneurons were labeled in A6 and A7 fields, as well as in the neighboring periventricular gray matter. In the midbrain, the IC was labeled intensely, as well as the substantia nigra. Finally labeled neurons have been identified in the ventrolateral part of the periaqueductal gray (PAG), and in the nuclei of the oculomotor complex.

### 3.6 Orexin-A

In the 9 years old specimen, in the medulla, OX-A axonal terminal fields were identified in the hypoglossal nucleus (XII), SOL anteroventrally (see Fig. 21), dorsal raphe (DR), the reticulotegmental nucleus (RtTg), the reticulotegmental nucleus lateral part (RtTgL), and the ventral gigantocellular reticular nucleus (GiCRt). In the pons, OX-A axon terminals in which there are many boutons were observed surrounding neurons of the LC in both the 9 years and 54 years specimens. In the 25 GW specimen, OX-A terminals were identified in the DR.

**Figure 21.**
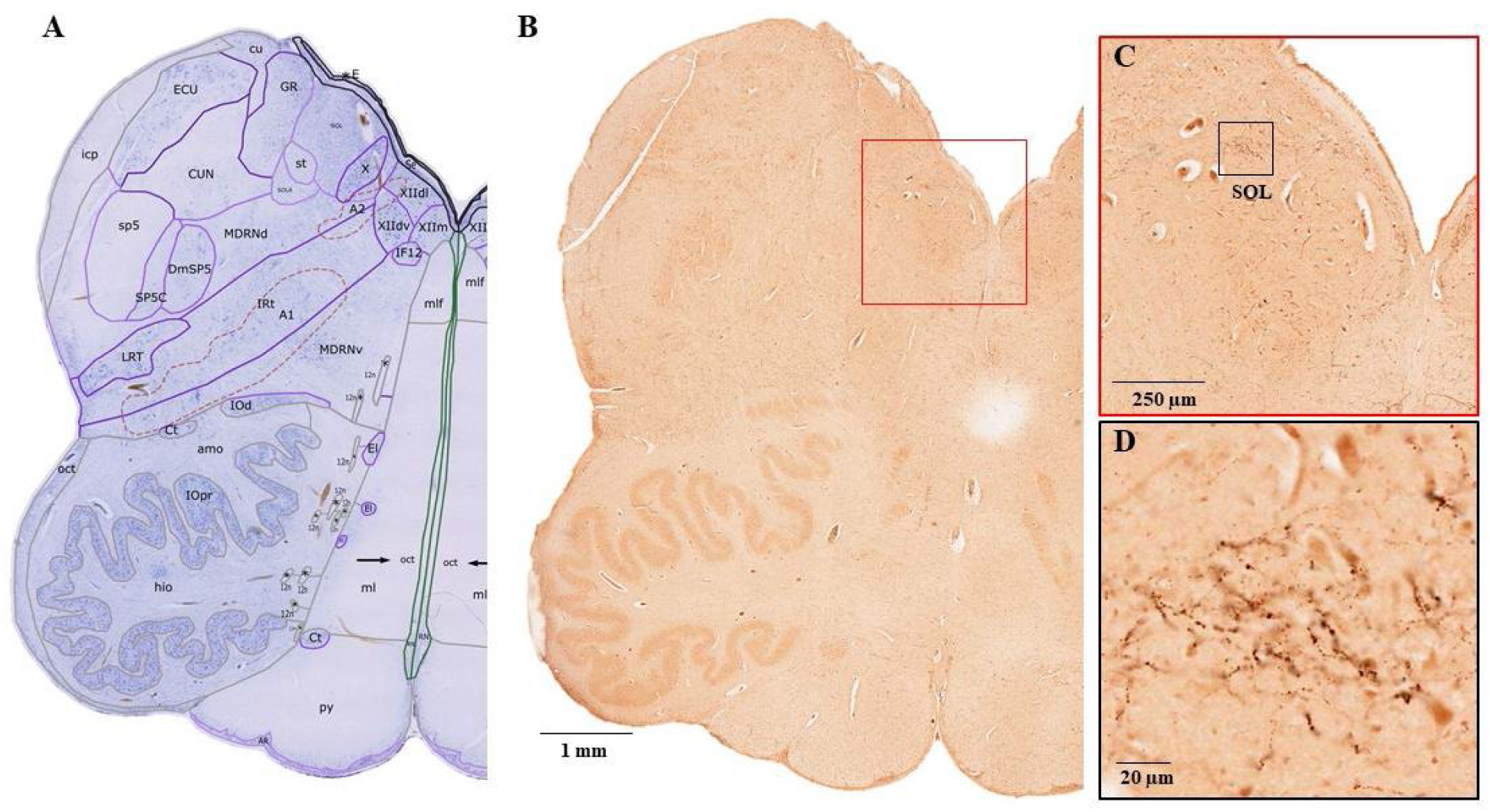
OX-A labeled axonal terminal fields in SOL. A. Nissl stained and annotated section at the level of SOL (obex: +4.0 mm). B. Corresponding OX-A stained section (obex: +3.88 mm). C and D. Details of OX-A labeled terminals and punctae in SOL.

### 3.7 Neurofilament heavy chain

NFH immunostaining revealed many neurons in a substantial number of nuclei in the brainstem in all three specimens. In the medulla, we identified the previously unreported protoplasmic commissural dendrites of the medial part of the hypoglossal nucleus (pcd; see Fig. 22) in the 9 year old specimen. These dendrites formed a distinct “commissure” between the hypoglossal nuclei.

**Figure 22.**
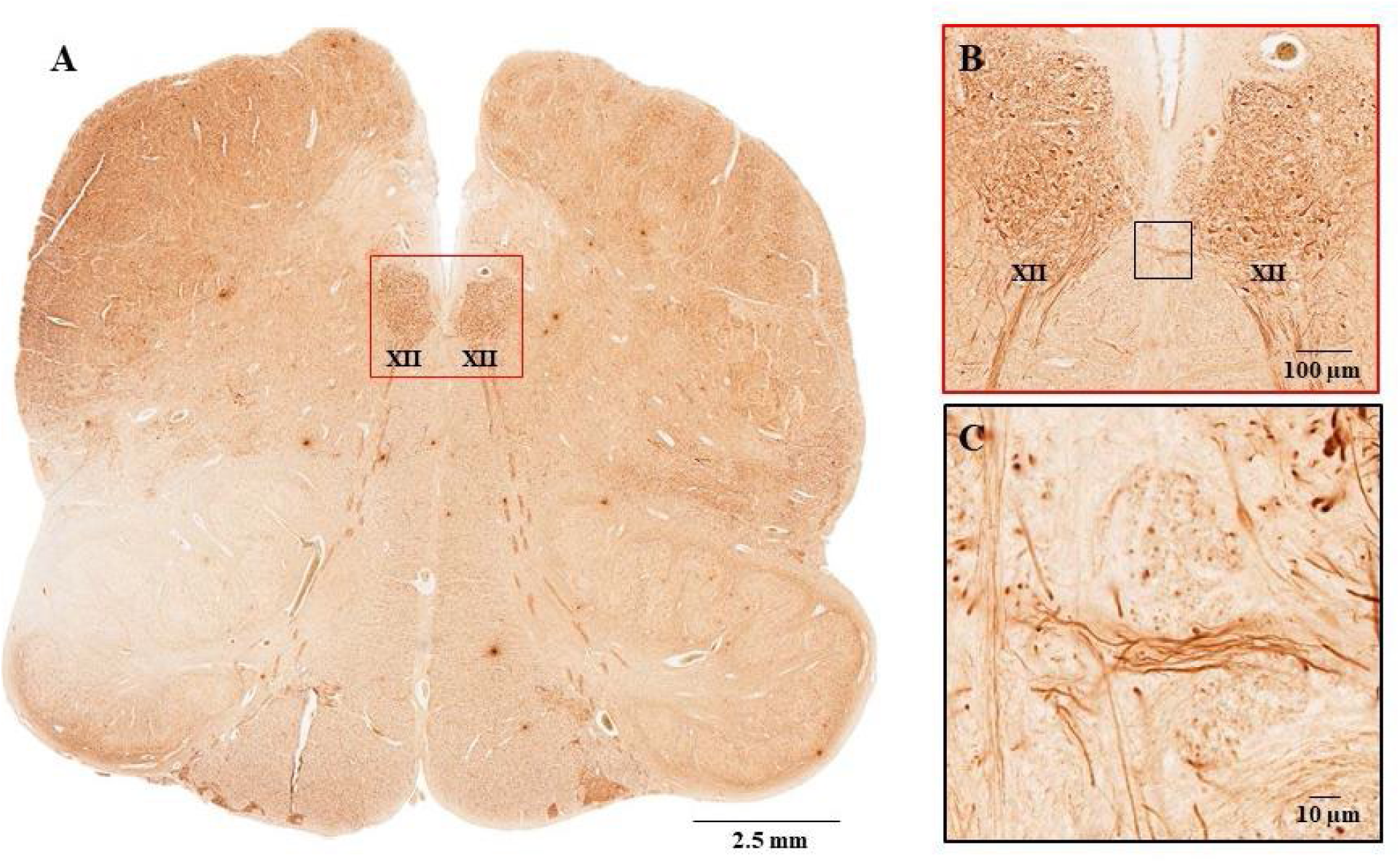
NFH labeled pcd in the hypoglossal nucleus, in the 9 years old specimen. A. NFH-stained section at the level of the hypoglossal nucleus, medial part (+1.04 mm). B and C. Details of A, showing the commissural dendrites of the hypoglossal nucleus, medial part.

In the pons, both neurons and fibres of the facial nerve nucleus were stained intensely (see Fig. 23). The NFH immunostaining of the facial nucleus allow us to further subdivide it in three parts.

**Figure 23.**
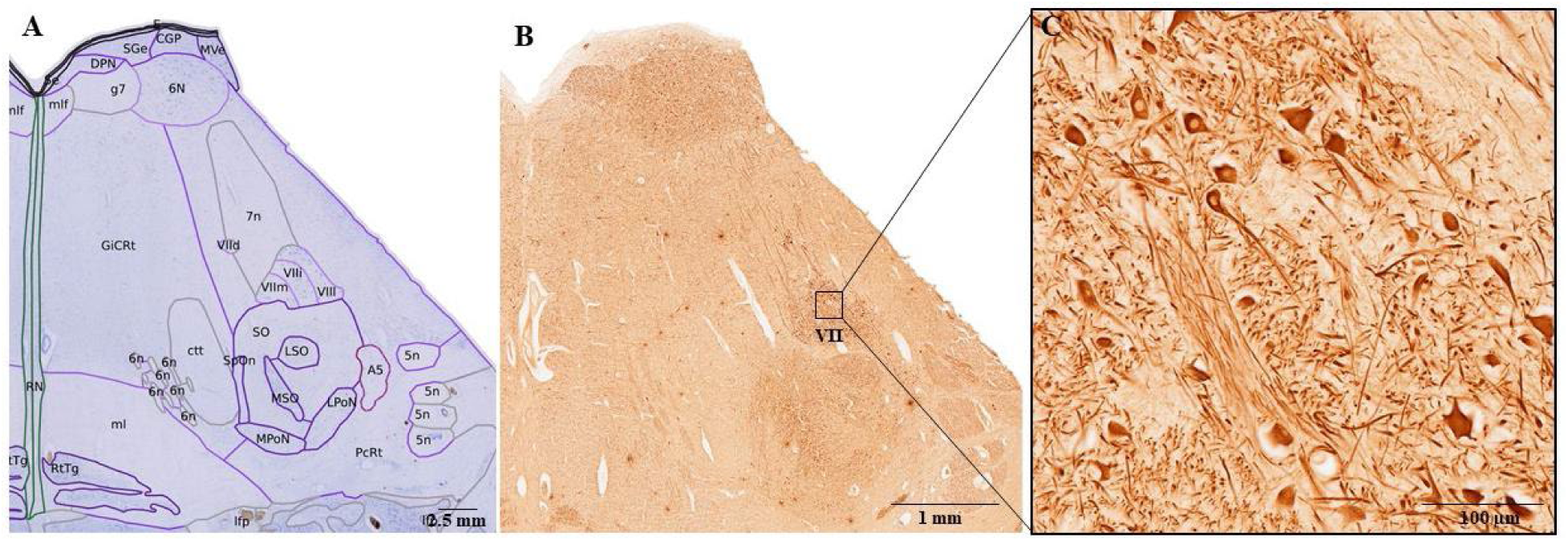
NFH labeled facial nucleus in the 9 years old specimen. A. NFH-stained section at the level of the facial nucleus (+14.4 mm). B and C. Details of A, showing labeled cells and terminals in the facial nucleus.

### 3.8 Pretectum

The pretectum (PRT) is located in the rostrodorsal part of the midbrain, bordered rostrally and dorsally by the medial and the lateral habenula, and the habenulo-interpeduncular tract. Medially it is bordered by the periaqueductal gray (PAG), and latero-rostrally by the thalamic posterior limitans (pli). Caudally and dorsolaterally it is bordered by the superior colliculus (SC), and ventrally by the midbrain reticular formation. The nuclei that are included in PRT are involved in processing of visual information (Gamlin, 2006), with the possible exception of the anterior pretectal nucleus (APT), which is involved in somatosensory and pain processing (Rees & Roberts, 1993). The nomenclature of the vertebrate pretectal nuclei has varied over time (Kuhlenbeck & Miller, 1949; Scalia, 1972; Hutchins & Weber, 1985; Borostyánkői-Baldauf & Herczeg, 2002). The PRT is considered to include at least five nuclei: the anterior pretectal nucleus (APT), the nucleus of the optic tract (NOT), the medial pretectal nucleus (MPT), the olivary pretectal nucleus (OPT), and the posterior pretectal nucleus (PPT; Gamlin, 2006). The APT can be subdivided in a compact (APTc) and a diffuse part (APTd; Scalia, 1972). Additionally, the ventral part of the PPT was identified in humans and cats as a distinct nucleus and named the central tegmental area (CTA; Borostyánkői-Baldauf et al., 1999; Borostyánkői-Baldauf & Herczeg, 2002). The APT is the most rostral and lateral nucleus of PRT. It has an elongated shape, is bordered laterally by pli and in the horizontal plane is located between the nucleus limitans of the thalamus and the OPT (see Fig. 24; Hutchins & Weber, 1985).

**Figure 24.**
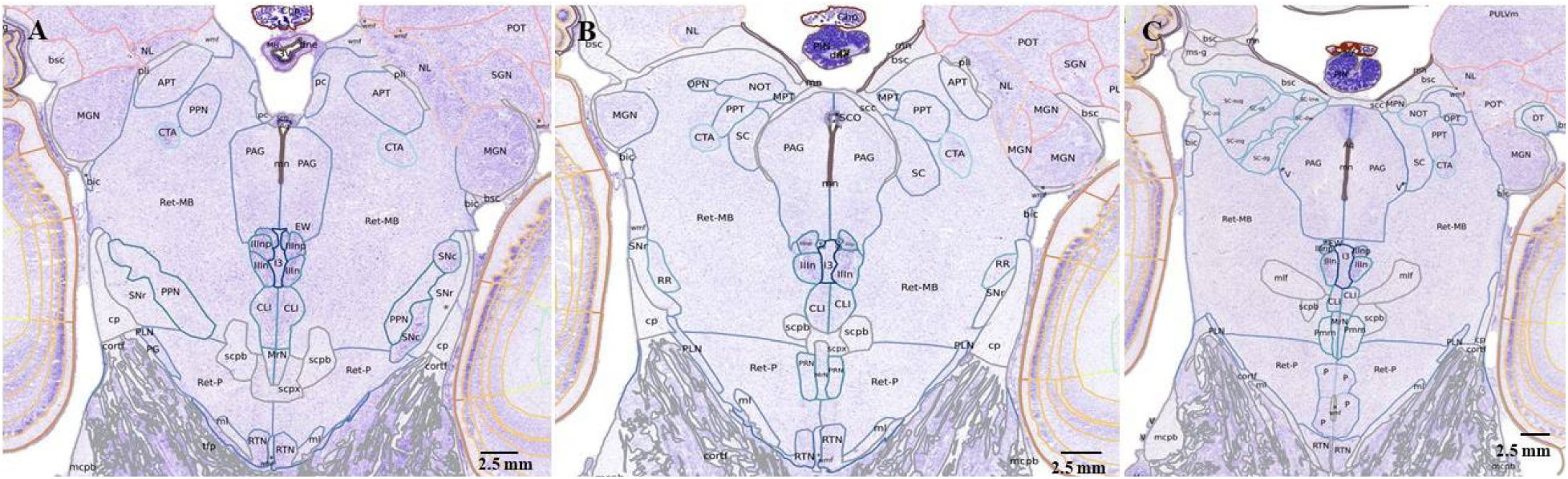
Nissl stained and annotated sections in the brainstem of the 25 GW specimen, showing the pretectal nuclei, anteriorly (A), mid antero-posteriorly (B), and posteriorly, at the level of SC (C).

The PPT is located ventromedially from APT, and lateral to PAG (Kanaseki & Sprague, 1974; Borostyánkői-Baldauf & Herczeg, 2002). In the 25 GW fetal brain, it starts at the posterior half level of APT and extends antero-posteriorly to the SC (see Fig. 24 A, B). The Nissl labeled neurons in PPT are medium to large, having different shapes, and are more prominent in the lateral and posterior parts of the nucleus.

The CTA is located ventrally from the PPT and in Nissl stain it appears as a rounded-triangular nucleus, labeled more intensely than PPT (see Fig. 24).

The MPT is the smallest nucleus within pretectal complex, and it is located lateral to the base of the pineal body (see Fig. 24 B, C). It extends from the caudal portion of habenulo-interpeduncular tract and ends rostral to the fibers of posterior commissure (Kanaseki & Sprague, 1974). In the 25 GW fetal brain, it is located between the commissure of the superior colliculus (scc) and the NOT. Labeled cells tend to be fusiform and horizontally oriented.

The NOT borders are difficult to define because the nucleus is partially embedded with fibers of the brachium of the superior colliculus (bsc). NOT is bordered rostromedially by the lateral habenula, and caudally it is bordered by the OPN (Kuhlenbeck & Miller, 1949; Hutchins & Weber, 1985; Borostyánkői-Baldauf & Herczeg, 2002). In the 25 GW fetal brain, we identified this nucleus lateral to the MPT and bordered laterally by the OPT (see Fig. 24B, C). Cytoarchitectonically, NOT includes intensely labeled pairs of cells, as described in the literature (Hutchins & Weber, 1985).

The OPT is located in the center of the PRT, lateral to the MPT, below the bsc, and it extends posteriorly into SC (Borostyánkői-Baldauf & Herczeg, 2002; see Fig. 24B, C). It has an ellipsoid shape, and its characteristic cytoarchitecture patterns are “whirlpools” of medium sized cells in Nissl staining (Hutchins & Weber, 1985; Borostyánkői-Baldauf & Herczeg, 2002).

## 4. Discussion

In this paper we introduce ANCHOR, which includes more than 800 histological sections of the human brainstem at three stages: fetal 25 GW, 9 years old, and 54 years old. The histological sections have been serially stained for Nissl, and for seven immunomarkers. Out of the 800 stained sections, more than 200 have been manually annotated. Overall, more than 200 parts of the human brainstem have been identified and annotated in the three specimens. This, together with BFI and MRI images for each of the specimens, comprise, to our knowledge, the most comprehensive data set of the human brainstem. ANCHOR also includes a publicly available online viewer that allows users to navigate across sections and modalities.

Key findings include the identification of a distinct fasciculus of protoplasmic commissural dendrites of the medial part of the hypoglossal nucleus (pcd), previously only identified in the African wild dog (Chengetanai et al., 2025), and the distribution of the catecholaminergic nuclei in the human brainstem. This is the first time, to our knowledge, that the distinct fasciculus formed by the pcd has been identified in the human brain.

The catecholaminergic nuclei in the human brainstem have been described previously (Pearson, 1983; Arango et al., 1988, Blessing & Gai, 1997; Halliday et al., 2012), as well as in other species (Smeets & Gonzalez, 2000; Manger & Eschenko, 2021; Williams et al., 2022). The TH-immunostained nuclei in the human brainstem described here are qualitatively similar to those in the brainstem of other primates (Williams et al., 2022). Additionally, we describe the TH positive ascending fiber tract from the A7 to PBl. The ANCHOR annotations of the catecholaminergic fields in the human brainstem across three stages, which complement the Nissl-based annotations, may provide a future reference for further studies. Finally, we describe the pretectal nuclei in the 25 GW fetal brainstem, for the first time, to our knowledge (Verma et al., 2025).

Future developments of ANCHOR will include extended sets of IHC stains, especially acetylcholine esterase and serotonin, in situ and transcriptomics maps.

**Appendix Table 1.**
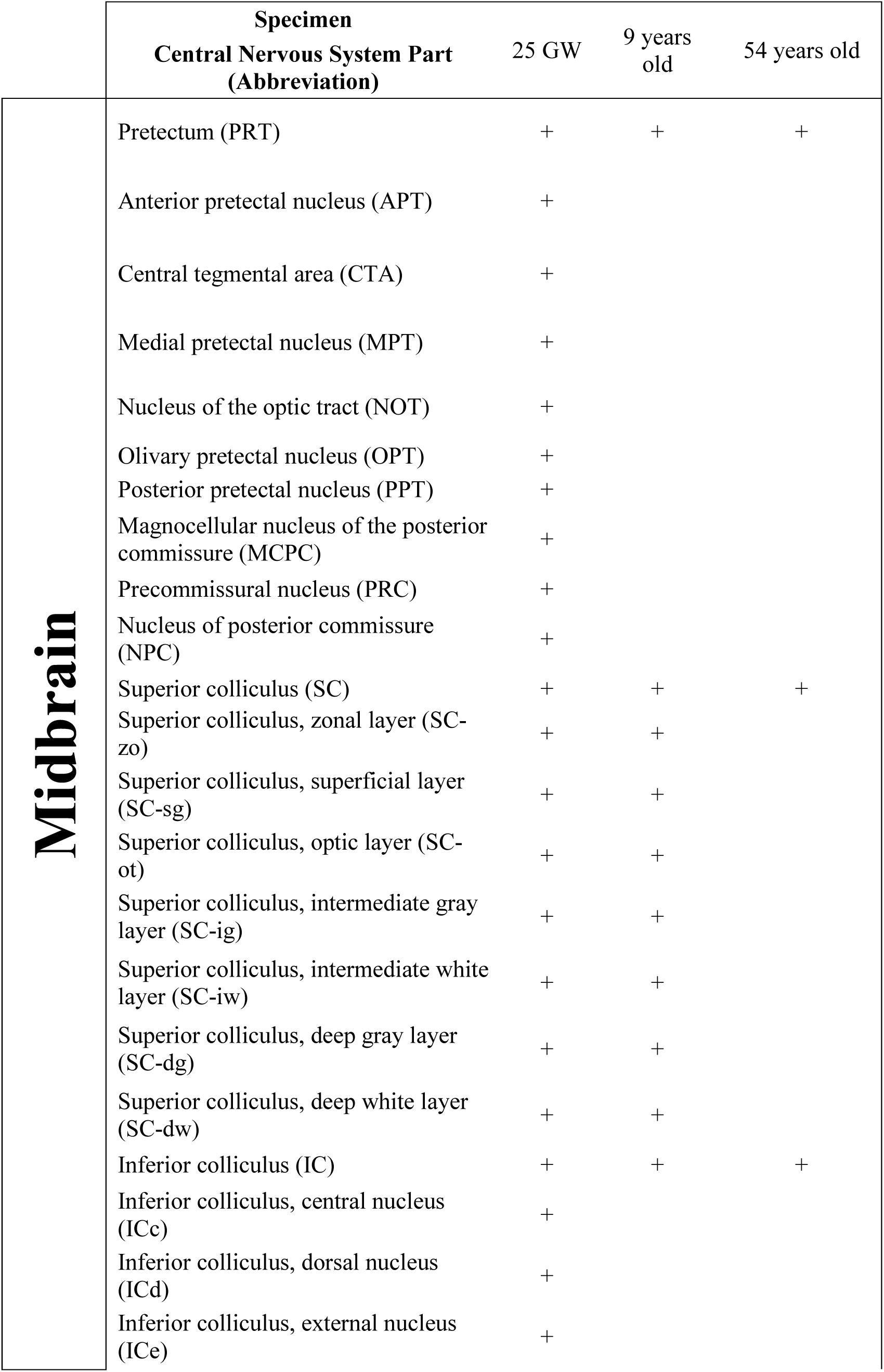
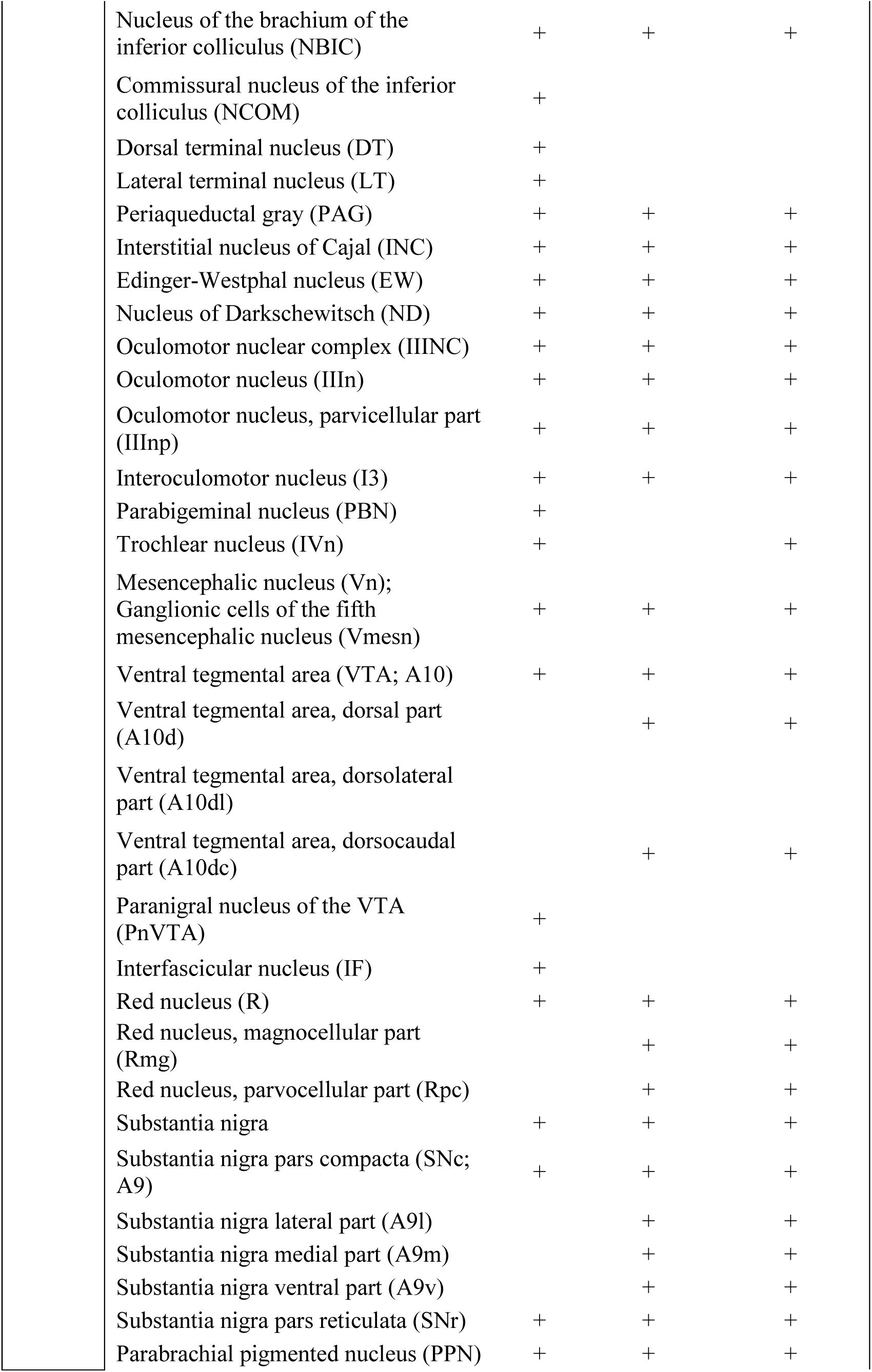
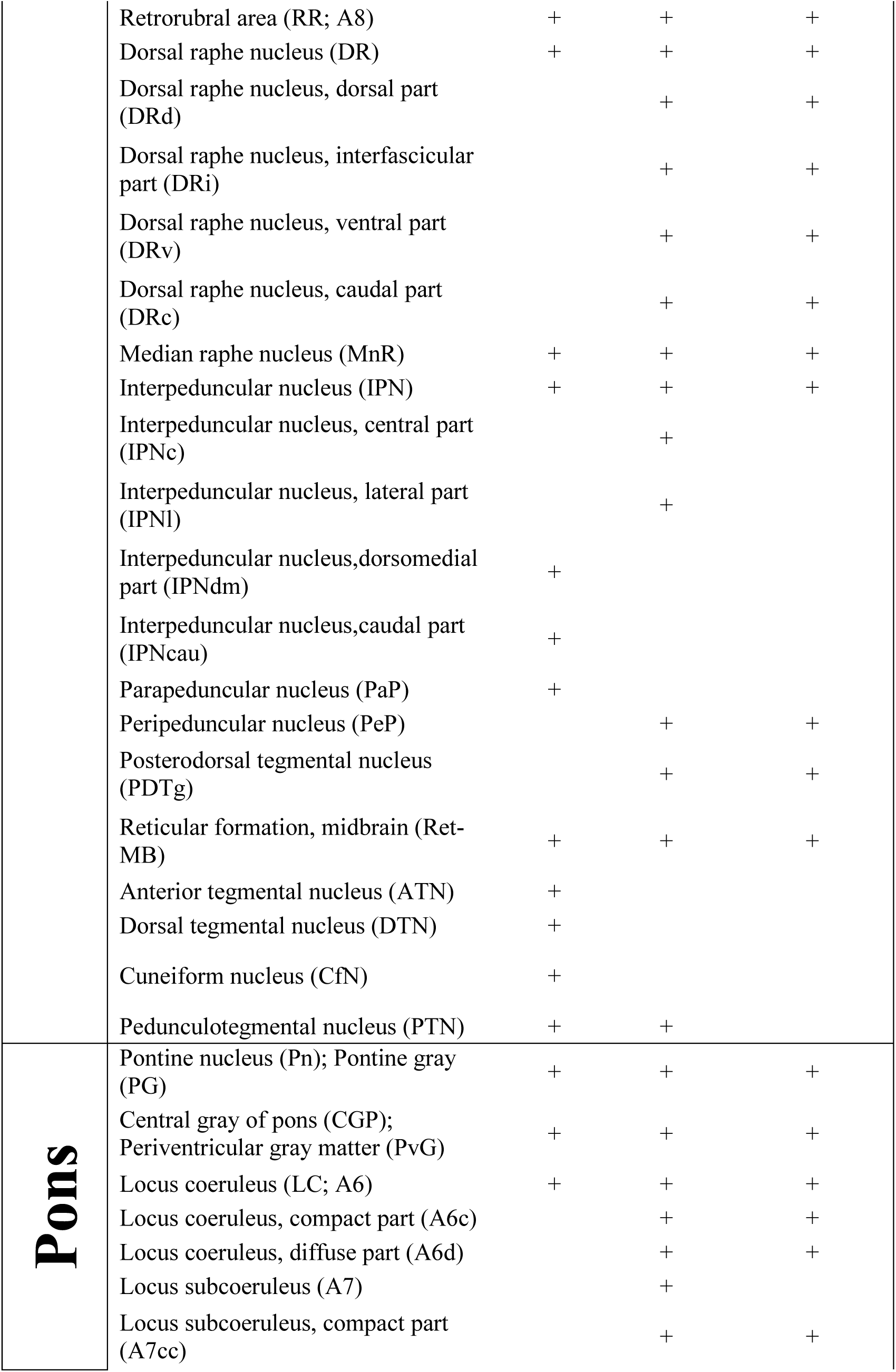
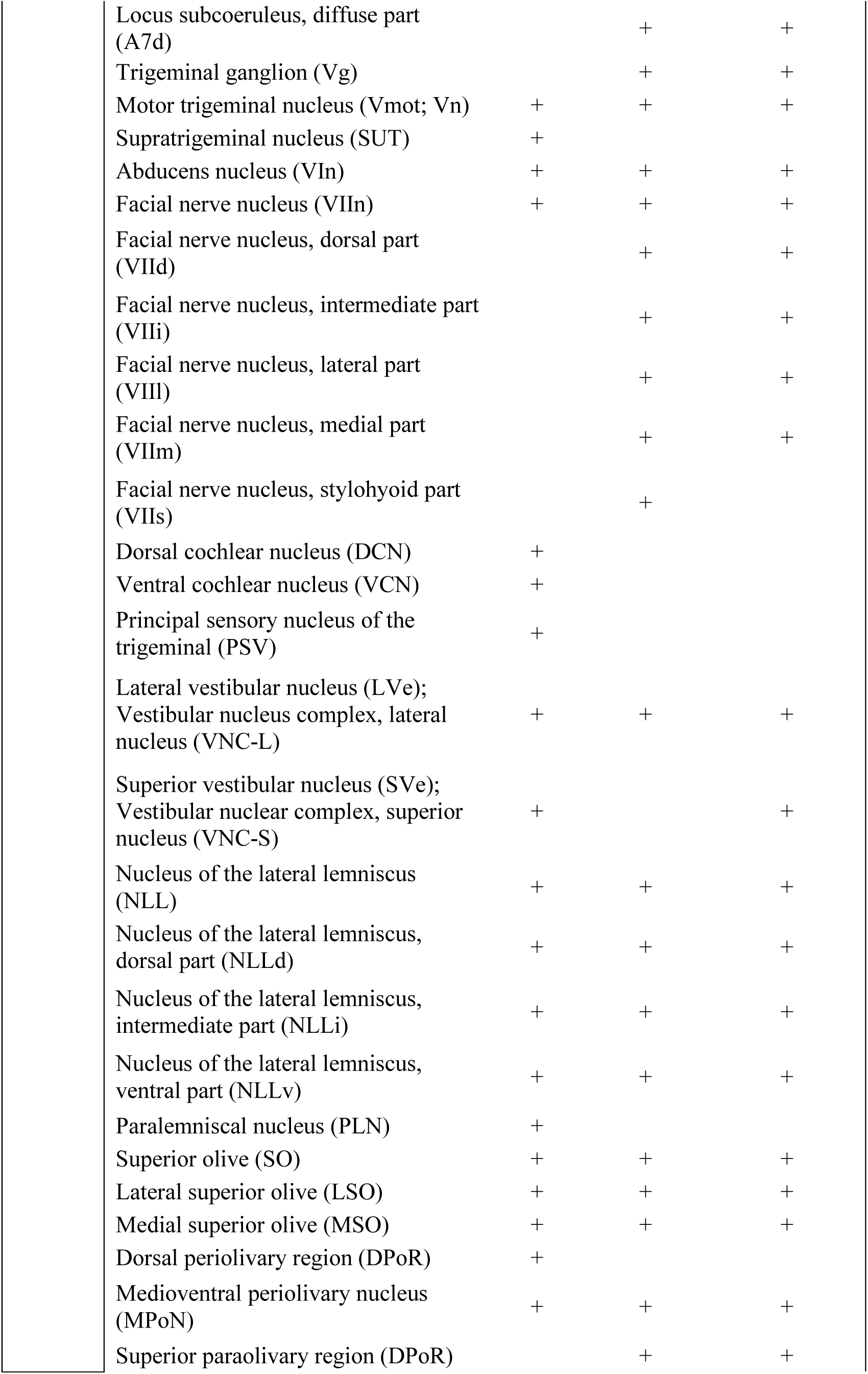
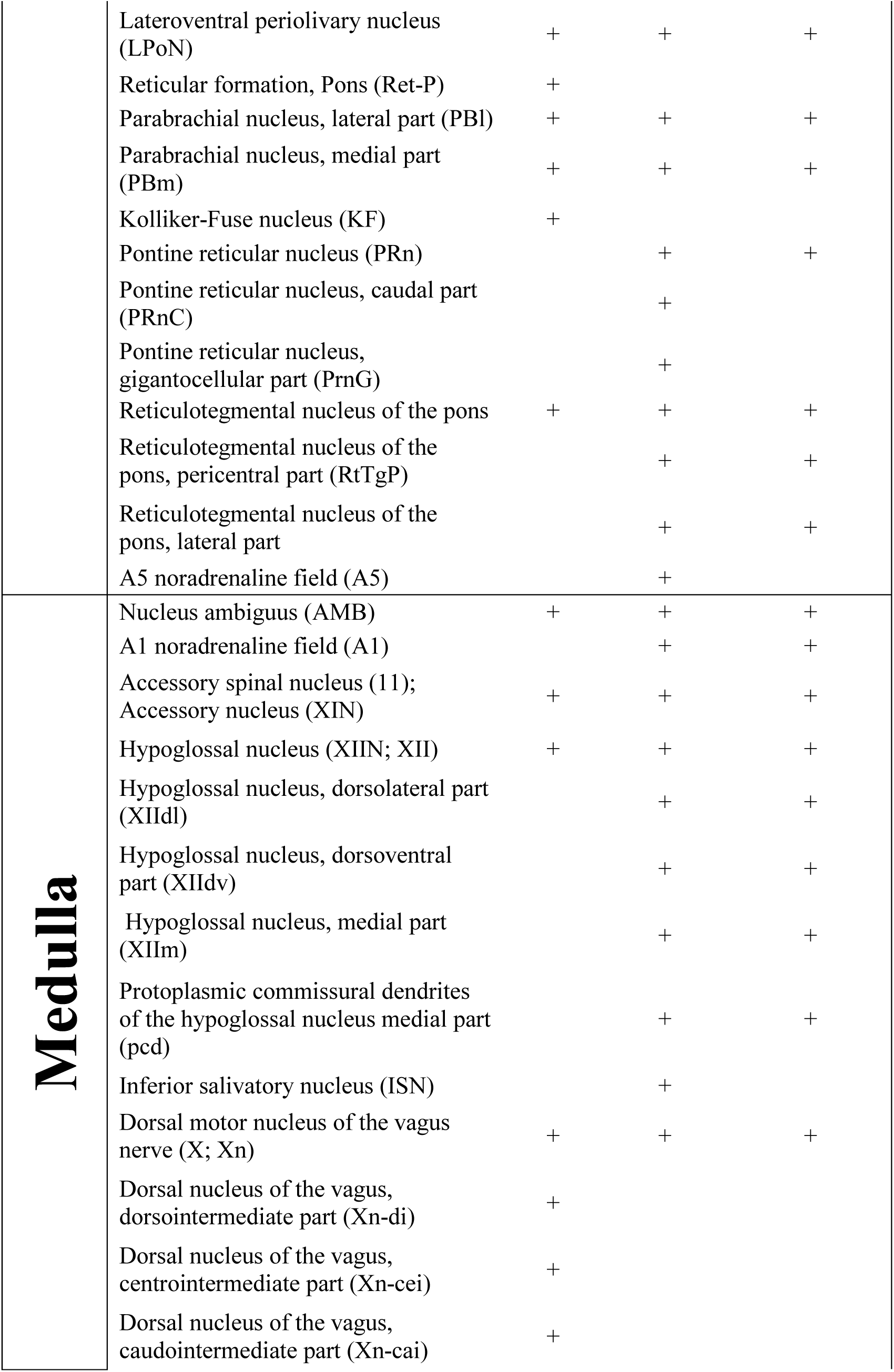
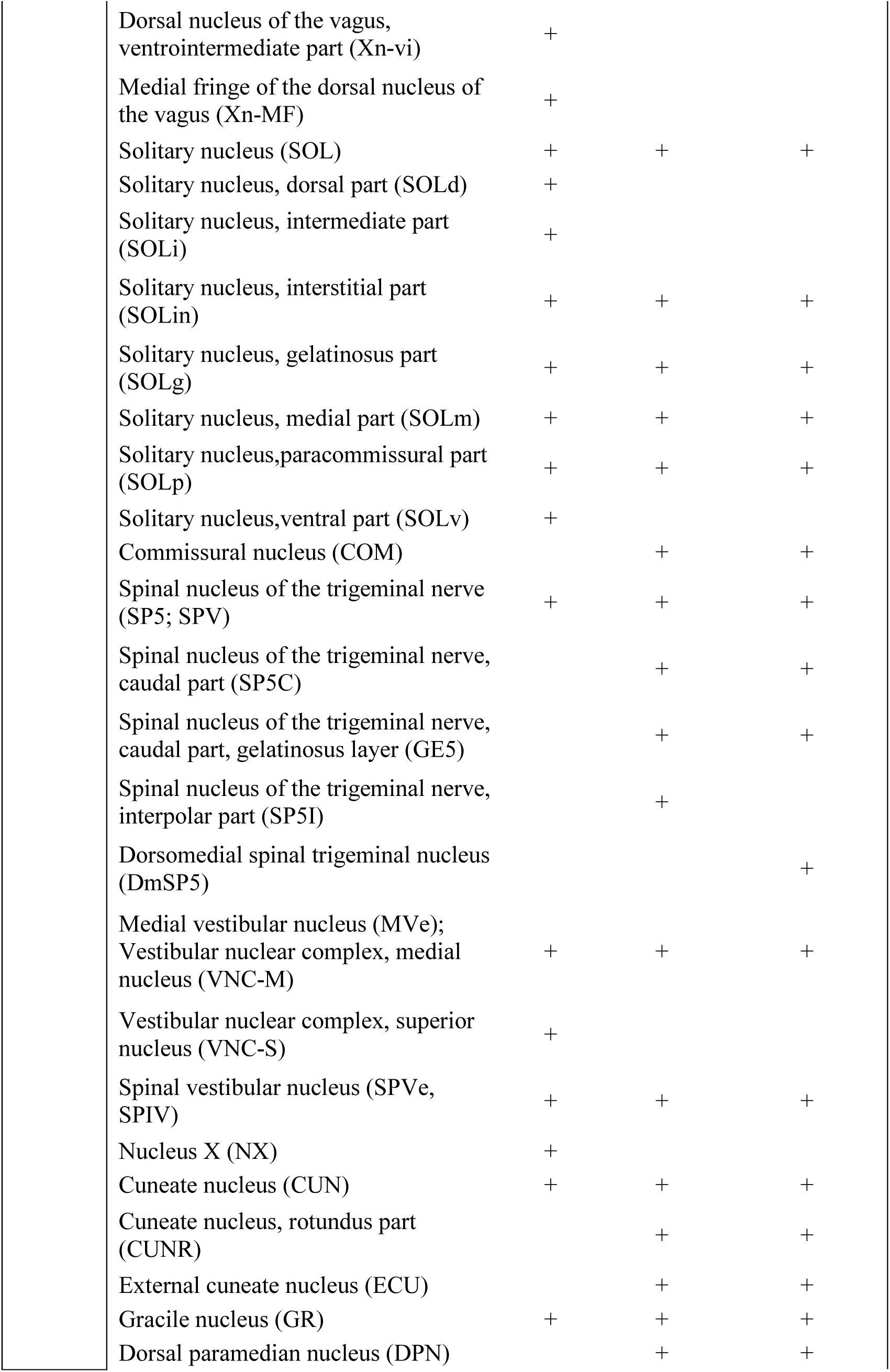
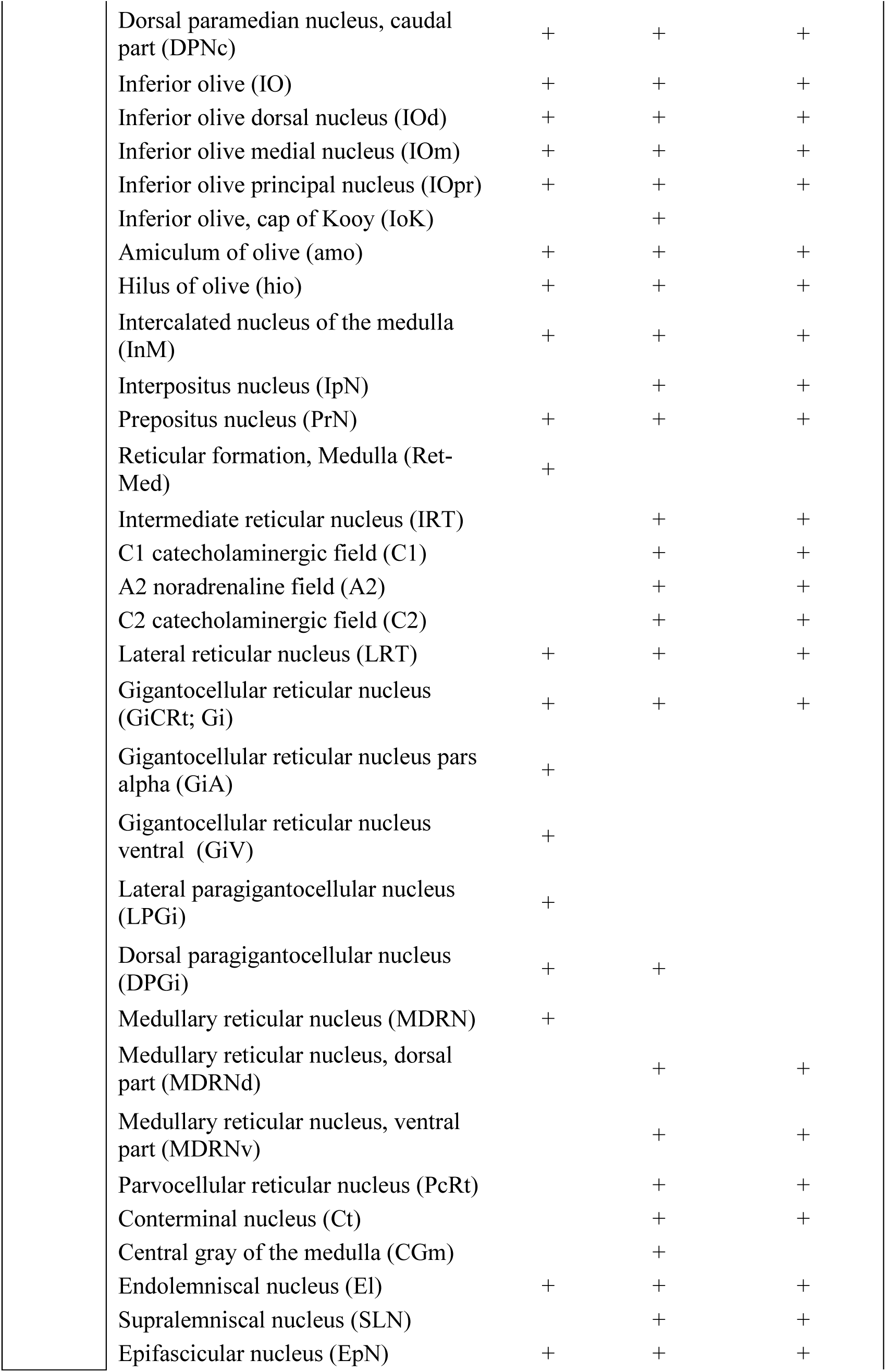
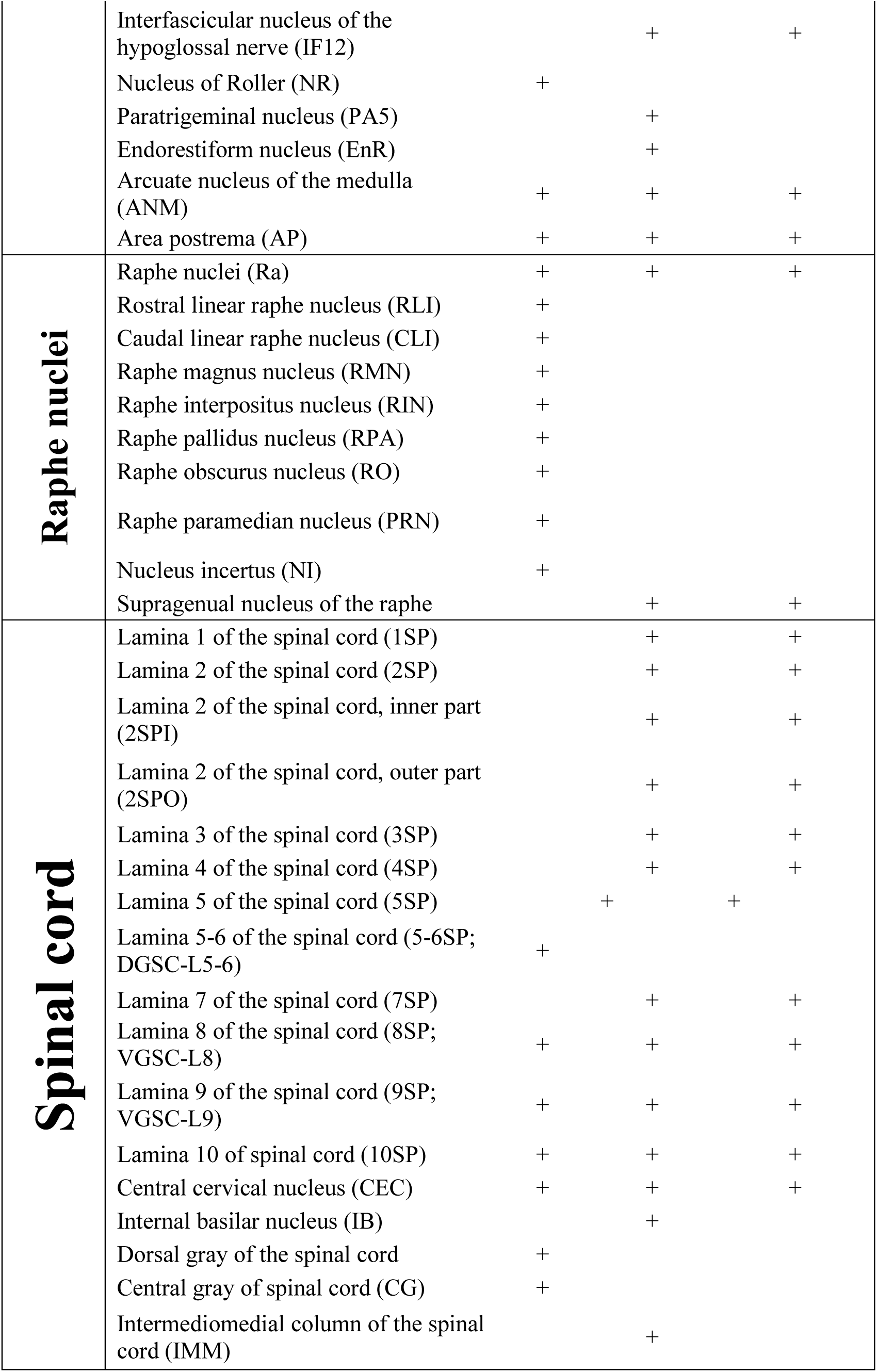
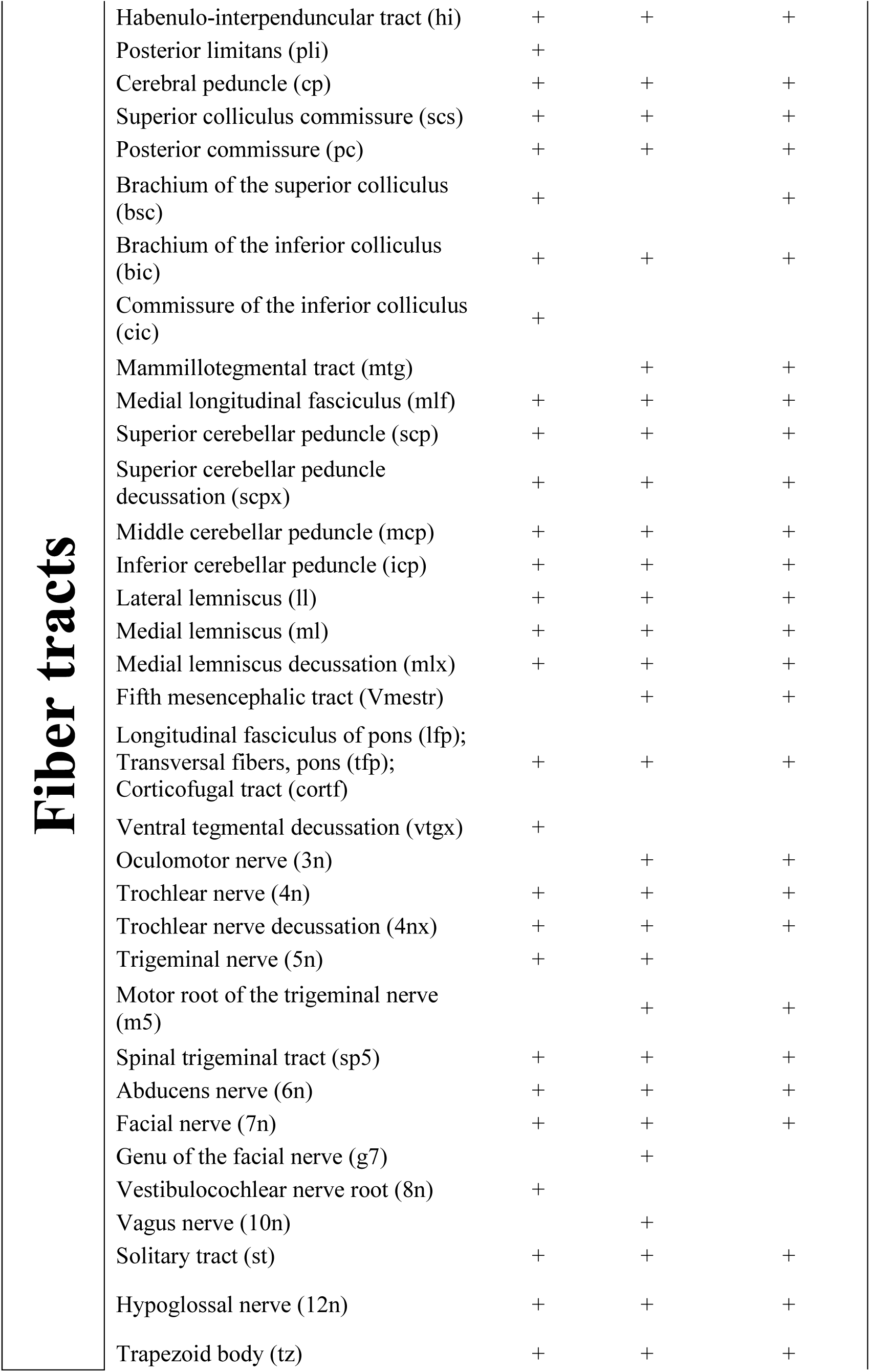
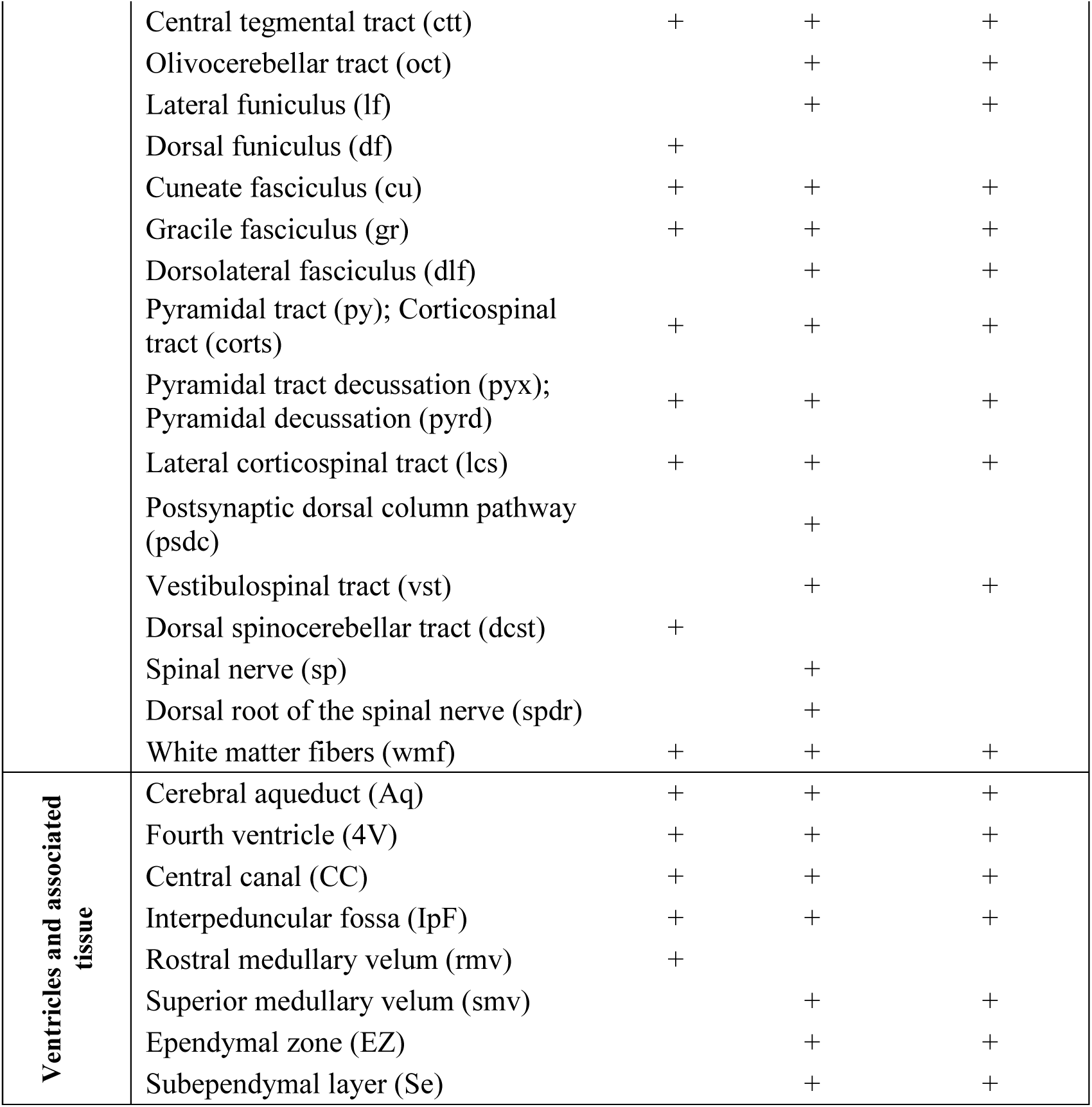
The nuclei, fiber tracts, ventricles and associated tissue, identified and annotated in all three specimens. Synonyms of terms are listed within the same cell.

